# Tissue-resident neutrophils serve homeostatic and immunological functions in embryos

**DOI:** 10.1101/2025.06.12.659311

**Authors:** Julian Hofmann, Laura Lintukorpi, Emmi Lokka, Venla Ojasalo, Sheyla Cisneros Montalvo, Damien Kaukonen, Lin Ma, Noora Kotaja, Heidi Gerke, Marko Salmi, Pia Rantakari

## Abstract

Development of neutrophils in the bone marrow and their crucial role in first-line defense are well understood in adults, but remarkably little is known about fetal neutrophils. Here, we analyzed the production, distribution, and functions of neutrophils during embryonic development in the mouse. We discovered that multiple non-hematopoietic steady-state organs harbor substantial numbers of immature and mature neutrophils, many of which are localized to tissue parenchyma outside the vessels. Using single-cell transcriptomic analyses, we revealed the presence of neutrophil progenitors and precursor cells in the blood and even in non-hematopoietic tissues in fetal and newborn mice. Embryonic tissue-resident neutrophils were transcriptionally different from embryonic blood-borne neutrophils and adult neutrophils. We demonstrated, through functional analyses, that embryonic neutrophils proliferated actively, had a high glycolytic capacity, and exhibited distinct diurnal rhythmicity. Embryonic neutrophils displayed lineage-specific innate immune effector functions and were responsive to maternal immunostimulation and immunosuppression. Using a genetic embryonic neutrophil depletion model, we discovered that neutrophils impact the piRNA pathway in the testis. Collectively, our data provides an atlas of the fetal neutrophil landscape and dissects their responses in steady-state.

**Summary:** Non-hematopoietic steady-state tissues harbor extravascular immature and mature neutrophils during fetal development. Embryonic neutrophils are endowed with multiple effector mechanisms and also serve homeostatic roles during tissue development.

## Introduction

Bone marrow-derived neutrophils are short-lived immune cells that migrate from the blood into sites of injury to engage in various antimicrobial functions, including the secretion of effector molecules and phagocytosis of pathogens (Koenderman et al., 2022; Ballesteros et al., 2020; Evrard et al., 2018; Aroca-Crevillén et al., 2024). In addition to their well-established role as the first line of innate immune defense in acute infections, studies in postnatal mice have revealed that neutrophils can infiltrate non-infected, steady-state tissues (Casanova-Acebes et al., 2018). In the bone marrow, spleen, and lung, a heavy neutrophil infiltrate was observed in the tissue parenchyma, whereas in tissues such as the liver, intestine, adipose tissue, skin, and muscle, only a few cells were present, typically located within the blood vessels. Interestingly, no neutrophils were observed in the brain, ovaries, and testis under steady-state conditions. The prolonged presence of neutrophils in many healthy tissues suggests that these myeloid cells may perform regulatory or supportive functions, possibly contributing to tissue homeostasis and immune surveillance (Puga et al., 2012).

In mice, neutrophils are first detected in the blood at embryonic day 11.5 (E11.5) (Wynn et al., 2017; da Silva Júnior et al., 2024). These early embryonic neutrophils differentiate from erythro-myeloid progenitor cells (EMPs) that colonize the fetal liver at E9.5, initiating the production of fetal erythrocytes, monocytes, and neutrophilic granulocytes. As the embryo develops, the granulopoiesis gradually shifts into bone marrow, where mature neutrophils are produced from hematopoietic stem cells via multipotential progenitors, granulocyte-macrophage progenitors, pro-neutrophils, pre-neutrophils, and immature neutrophils. However, the presence of neutrophils in fetal tissues (apart from tissues with hematopoietic activity) and their potential functions in an essentially sterile milieu in utero remain unexplored.

Macrophages, the other major phagocytic leukocyte type in the body, are distributed to different organs early during ontogeny and support multiple non-immunological functions (Park et al., 2022). Here, we propose that neutrophils may also infiltrate into non-hematopoietic tissues during the fetal period to support homeostatic functions in addition to serving as guardians against potential intrauterine microbial challenges. By combining in-depth single-cell RNA sequencing (scRNA-Seq) and imaging, we demonstrate that neutrophils are present early in most steady-state embryonic tissues and exhibit distinct transcriptional profiles compared to their adult counterparts. Functional assays revealed that embryonic neutrophils are already capable of performing essential neutrophil functions, such as reactive oxygen species (ROS) production and phagocytosis. Notably, we found that embryonic neutrophils support P-element induced wimpy interacting (PIWI) RNA (piRNA) metabolism and DNA methylation during testicular development, suggesting that they also serve nonimmune regulatory functions during the development of normal tissues.

## Results

### Immature and mature neutrophil populations are present in non-hematopoietic tissues in embryos

To characterize steady-state neutrophils during the development, we performed flow cytometric analyses from multiple tissues of wild-type (WT) mice housed under specific pathogen-free conditions by analyzing Ly-6G expression in myeloid cells. In newborn (NB) mice, we observed neutrophils identified as CD11b^+^Ly-6G^+^ cells not only in the hematopoietic organs (liver and spleen), but also in the lungs, pancreas, kidneys, and testes. On the other hand, in the thymus and brain, no or only very occasional CD11b^+^Ly-6G^+^ cells were found (Fig. S1A-C). To determine the kinetics of neutrophil infiltration in detail, we collected blood and tissues from steady-state mice at embryonic day (E) 17.5, at birth, and postnatally at 1, 2, 5, and 8 weeks of age (Fig. 1A-B). In the liver, we found that 70-90% of the CD45^+^CD11b^+^SiglecF^neg/low^Ly-6C^+^Ly-6G^+^ cells (hereafter assigned as neutrophils) had an immature CD101^low^CXCR2^low^ (Evrard et al., 2018) phenotype in the prenatal, NB, and 1-week-old mice (Fig. S1D-F). In 2-week-old and older mice, in contrast, less than 30% of the liver neutrophils were immature (Fig. S1F). This transition coincided with a decrease in the frequency and absolute numbers of neutrophils in the liver at 2 weeks of age (Fig. S1E). In the bone marrow, neutrophils were present from E17.5 to 8-week-old mice (Fig. S1G-I), their absolute number increased exponentially with time, and the immature neutrophils dominated at each time point without any declining trend (Fig. S1H). In the blood, where the frequency of neutrophils peaked at birth, the proportions of immature and mature neutrophils were roughly equal from E17.5 to 1-week-old mice (Fig. S1J). In contrast, thereafter, the vast majority of the cells displayed the mature phenotype (Fig. S1K).

**Figure 1.**
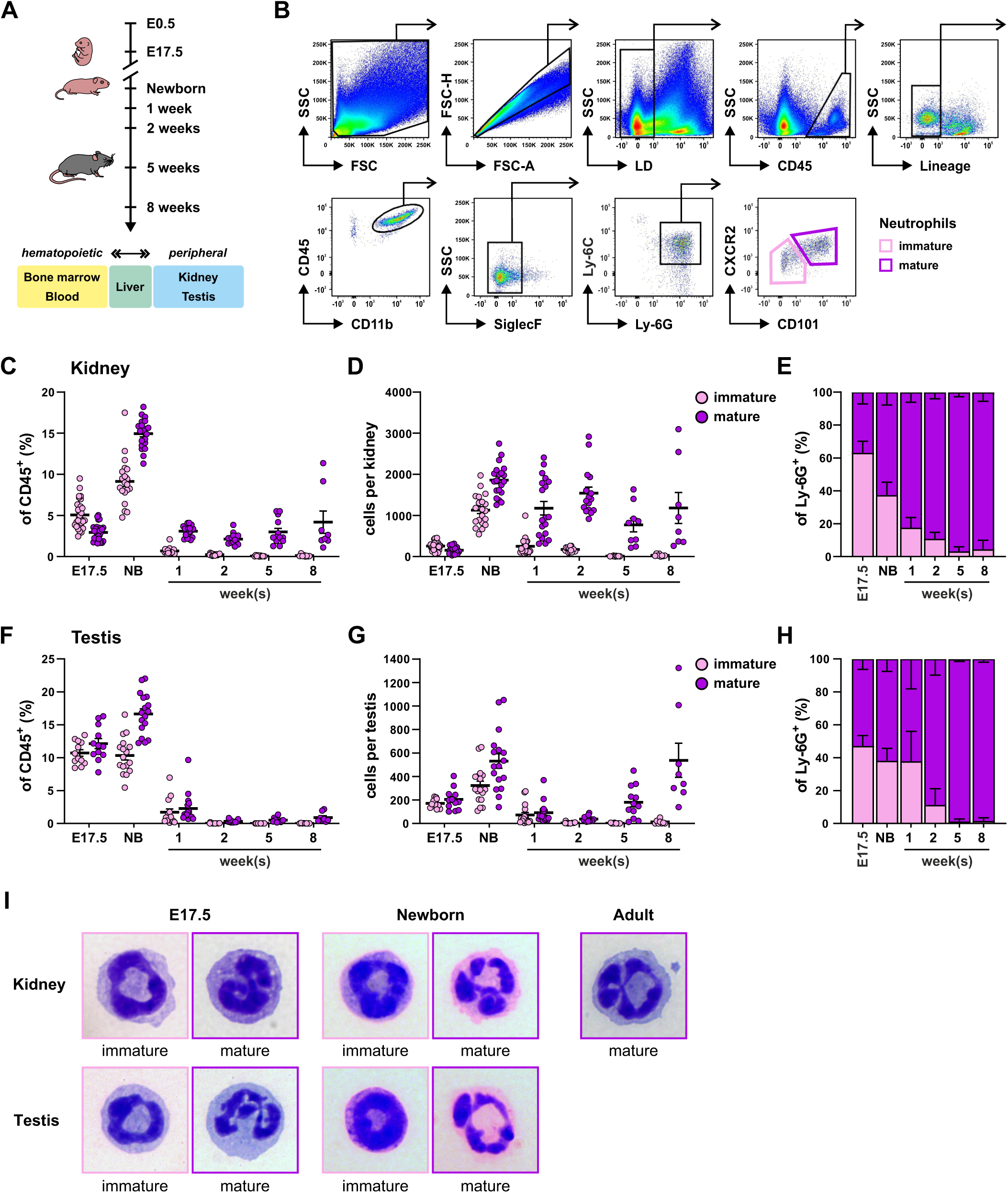
Immature and mature neutrophils are present in steady-state non-hematopoietic tissues in embryos. **(A)** Schematic experiment overview. **(B)** Flow cytometry gating strategy with representative plots from newborn kidney for immature CD101^-^CXCR2^-^ (pink) and mature CD101^+^CXCR2^+^ (purple) neutrophil populations. **(C-E)** Frequency (C), numbers (D), and proportions (E) of immature and mature neutrophils in the kidney at the indicated developmental time-points. **(F-H)** Frequency (F), numbers (G), and proportions (H) of immature and mature neutrophils in the testis at the indicated developmental time-points. **(I)** Representative cytological images of sorted, Wright-Giemsa-stained immature and mature neutrophils in the kidney and testis. In C and D, each dot represents both kidneys from one mouse (E17.5 n = 29, newborn; n = 20, 1 week; n = 19 and 2 weeks; n = 16 mice) or one kidney from each mouse (5 weeks; n = 9 and 8 weeks; n = 8). In F and G, each dot represents a pool of 4 testes from 2 mice (E17.5 n = 12 pools) or both testes of one mouse (newborn; n = 17, 1 week; n = 16 and 2 weeks; n = 16 mice) or one testis from each mouse (5 weeks; n = 12 and 8 weeks; n = 8). Data are presented as mean ± SEM (C, D, F, G) and as mean ± SD (E and H). All data are from 3-7 independent experiments.

We notably observed substantial numbers of neutrophils in non-hematopoietic tissues during early development (Fig. 1C-H and S1C). In the kidney, 5-10% of leukocytes were immature neutrophils and 3-15% mature neutrophils at E17.5 and in NB. The number of mature neutrophils was at the highest level (1000-2000 cells/kidney) in NB mice but remained detectable until the end of the 8-week analysis period (Fig. 1D). A similar pattern of a predominantly pre- and perinatal neutrophil infiltrate (20-30% of all leukocytes) consisting of both immature and mature cells was observed in the testis (Fig. 1F and G). Analyses of the cellular morphology of sorted Ly-6G^+^ cells revealed a characteristic band and ring-shaped multilobular nuclear morphology of immature and mature neutrophils, respectively, already in E17.5 and NB mice tissues (Figures 1I and S1L). Collectively, our data reveal the infiltration of immature and mature neutrophils in non-hematopoietic organs in steady-state developing mice, which is most prominent in fetuses and at birth.

### Embryonic neutrophils are extravascular in the tissue parenchyma

To define the distribution of neutrophils within embryonic tissues, we used a neutrophil-specific *Ly6G ^Cre-tdT/Cre-tdT^* (Hasenberg et al., 2015) reporter mouse and crossed it with *R26^tdT/tdT^* mice for improved tracking of neutrophils via tdTomato (tdT) expression (Fig. S2A). Our flow cytometry analyses revealed 25% to 35% labeling efficiency of Ly-6G^+^ neutrophils in different tissues of E17.5 neutrophil-specific reporter embryos (Fig. S2B). Using 3D confocal imaging of optically cleared whole mount tissues, the tdT^+^ neutrophils were detected in all hematopoietic tissues studied (liver, bone marrow, spleen; Fig. S2C-E). In the E17.5 kidney, neutrophils were scattered throughout the tissue parenchyma (Fig. 2A and S2F). Although a few neutrophils were found close to vessel walls, the majority of neutrophils were clearly outside the vessels (Fig. 2A and B). Occasionally, we found neutrophils in small clusters both in the cortex and outer medulla of the kidney (Fig. 2C and S2F). Occasionally, we also observed embryonic neutrophils lodged between tubule epithelial cells (Fig. S2G). In the E17.5 testis, neutrophils were concentrated around the highly vascular connective tissue in the mediastinum and the rete testis area that is involved in the transport of sperm cells from the testicle to the epididymis (Fig. 2D and S2H) (Kulibin and Malolina, 2020). Compared to the kidney, neutrophils in the testis were more frequently observed close to the outer surface of blood vessels (Fig. 2E-G), but only a few cells were found inside vessels (Fig. 2F).

**Figure 2.**
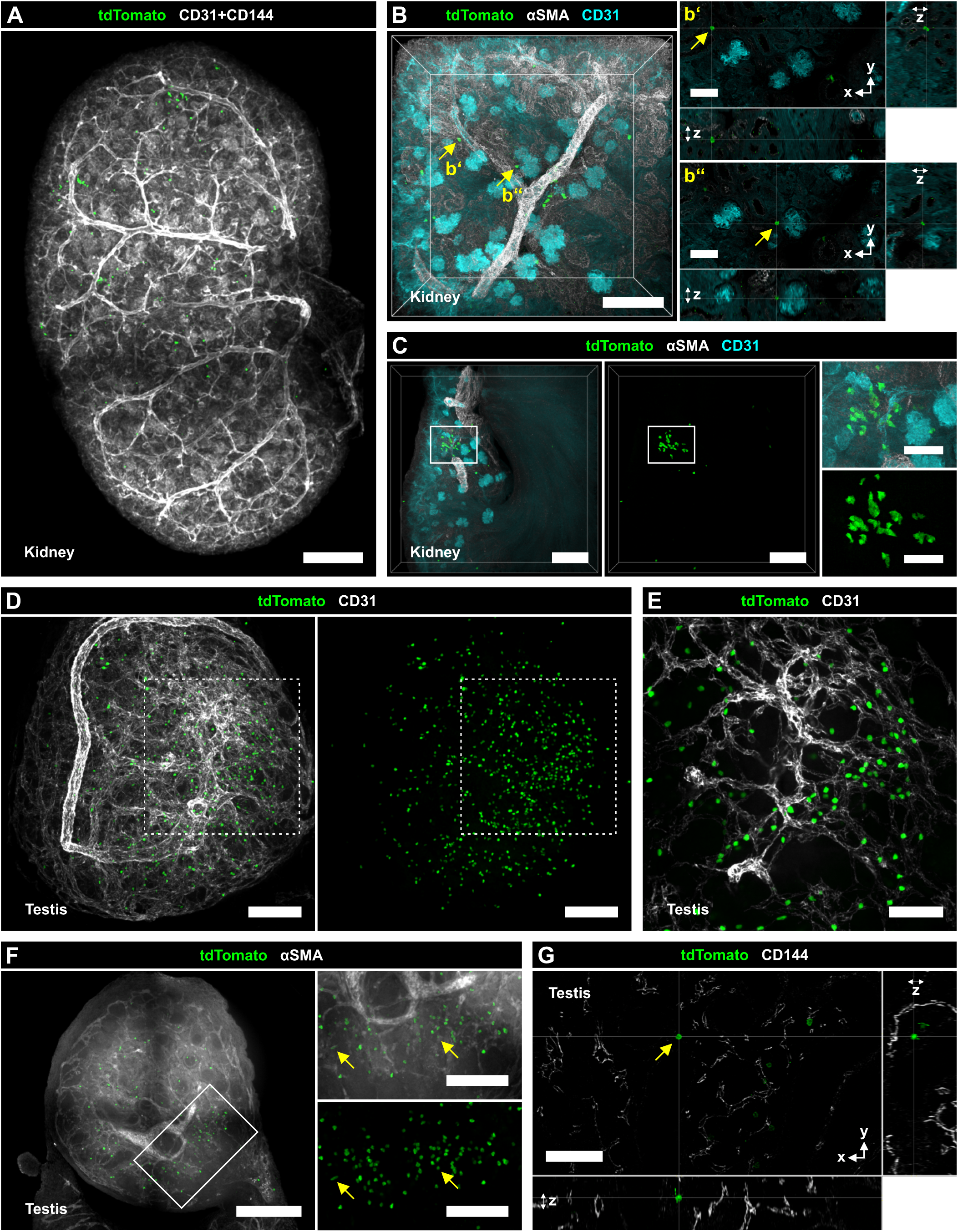
Embryonic neutrophils are largely extravascular in steady-state non-hematopoietic tissues. Representative confocal immunofluorescence microscopy images from at least three independent experiments of optically cleared wholemount kidney and testis from embryonic (E17.5) neutrophil reporter mice showing tdTomato-labelled (tdT^+^) neutrophils (green) and labelled vasculature (with the indicated antibodies; gray, cyan) as indicated. **(A)** Overview image of the kidney. Scale bar 200 µm. The same image has been used in Fig. S2F. **(B)** Cellular resolution volumetric image of the kidney with magnified cross-sectional views (**b’ and b’’**) of representative extravascular tdT^+^ cells (arrows). Scale bars 150 µm (full), 50 µm (magnified). **(C)** Imaging of the cortex area of the kidney with magnified views of the highlighted region (box). Scale bars 150 µm (full), 50 µm (magnified). **(D)** An overview image of the testis. Scale bars 100 µm. **(E)** Higher resolution image of mediastinum and the rete testis area highlighted in **D** (dashed box). Scale bar 50 µm. **(F)** Volumetric cross-section of testis and connected epididymis and magnified views of the mediastinum and the rete testis area (box) highlighting representative intravascular tdT^+^ cells (arrows). Scale bars 200 µm (full), 100 µm (magnified). **(G)** Magnified cross-sectional views of representative extravascular tdT^+^ neutrophil highlighted in Fig. S2H (arrow). Scale bar 50 µm. Different perspectives of the same images have been used in Fig. S2H.

To better discriminate the tissue-resident and intravascular neutrophils, we performed intravascular administration of fluorescently labeled Ly-6G antibody into the umbilical veins of embryos at E17.5. When 95-100% of blood neutrophils were labeled, we detected both Ly-6G-labeled (indicative of intravascular neutrophils) and unlabeled cells (indicative of extravascular) in varying proportions in flow cytometric analyses of the kidney and the testis (Fig. S2I), which is in line with our imaging data. Altogether, our observations revealed that a significant extravascular neutrophil population exists in the parenchyma of non-hematopoietic tissues in normal embryos.

### Embryonic neutrophils show a linear developmental trajectory

Next, we utilized scRNA-seq to explore the characteristics of embryonic neutrophils during steady-state tissue development (Fig. 3A and S3A). We sorted and sequenced CD45^+^ cells from the bone marrow, liver, blood, kidney, and testis of E17.5 and NB WT mice. To compare neutrophils from embryos and adults, we further integrated our embryonic data with published scRNA-seq data of sorted leukocytes from blood and bone marrow of normal 8- to 15-week-old mice (Ballesteros et al., 2020; Xie et al., 2020; Evrard et al., 2018)(Fig. S3A). Leukocyte progenitors (containing a mixed population of monocytic, neutrophilic, eosinophilic, and basophilic progenitor subsets) and differentiated leukocyte subpopulations were identified based on known marker genes (Fig. S3B and C).

**Figure 3.**
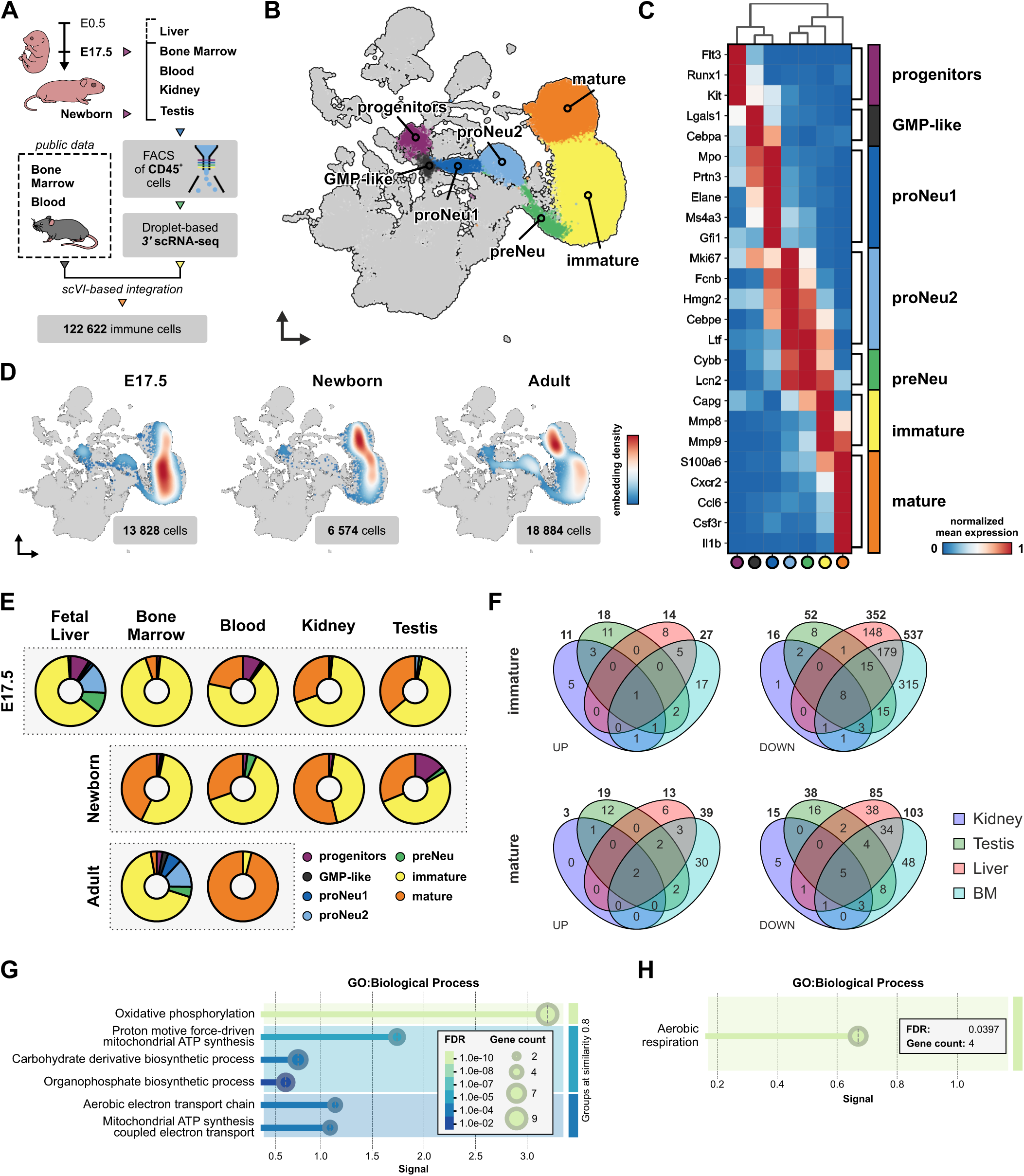
Embryonic neutrophils show a linear development pathway and tissue-selective gene transcription. **(A)** Schematic outline of sample collection and processing for scRNA-sequencing from steady-state tissues. **(B)** Uniform Manifold Approximation and Projection (UMAP) of the integrated scRNA-sequencing dataset. The different neutrophil maturation stages are indicated. **(C)** Matrixplot of normalized expression for selected marker genes across neutrophil subpopulations. **(D)** UMAP embedding density (cell distribution heatmap) and cell numbers of neutrophil ontogeny by age point. **(E)** Frequency plots of the neutrophil subpopulations by age and tissue of origin. **(F)** Venn diagrams showing the number of genes differentially upregulated (left) or downregulated (right) in immature (top) or mature (bottom) neutrophils from E17.5 kidney, testis, liver, and bone marrow compared to cells isolated from blood. **(G and H)** STRING functional enrichment visualization top 3 groups of gene ontology (GO) terms related to biological processes enriched in genes significantly upregulated in immature E17.5 neutrophils of bone marrow (G) and liver (H) compared to blood. FACS, fluorescence-activated cell sorting, BM, bone marrow; FDR, false discover rate corrected p-values; signal, weighted harmonic mean between the observed/expected ratio and -log(FDR).

To focus on neutrophils, we performed fine-resolution clustering and annotation of progenitors, committed proliferative neutrophil progenitors and precursors (preNeu) (Evrard et al., 2018), and immature and mature neutrophils (Fig. 3B and S3B-C and Data S1). Unbiased analysis of differentially expressed genes across seven clusters identified a common hematopoietic progenitor cluster expressing genes such as *Flt3, Runx*1, and *Kit* and a cluster of granulocyte-monocyte progenitor-like cells (GMP-like) marked by the expression of *Cebpa* (Fig. 3C and S3D and E). Two clusters of pro-neutrophils (proNeu1 and proNeu2) and a pre-neutrophil (preNeu) cluster were also detectable (Evrard et al., 2018). Finally, the biggest neutrophil cluster expressed markers of immature neutrophils (e.g., *Mmp8* and *Mmp9*), and the seventh neutrophil cluster represented mature neutrophils expressing, e.g., *Cxcr2*, and *Il1b* (Fig. 3C and S3D and E). The pseudotime trajectory analysis projected the clusters onto a single continuum developmental trajectory (Fig. S3F). In vivo validation using 5-bromo-20-deoxyuridine (BrdU) labeling, which marks proliferating cells and enables tracking of their differentiation (Evrard et al., 2018; Yona et al., 2013), revealed that only immature neutrophils were present between the 2- and 24-hour time points in the fetal liver. Immature neutrophils differentiated into mature forms by the 48-hour time point and had largely disappeared after 72 hours. Concurrently, the number of mature neutrophils increased (Fig. S3G), suggesting conserved developmental dynamics between embryonic and adult neutrophils (Evrard et al., 2018).

Similar to adult bone marrow, distinct neutrophil populations were also observed at the E17.5 and newborn tissues (Fig. 3D). However, E17.5 mice had the highest frequency of early progenitors, while the proportion of mature neutrophils was the lowest at E17.5 when compared to newborn and adult mice (Fig. 3D and E). We found distinct cluster-to-tissue distribution patterns among neutrophil sub-clusters (Fig. 3E). In the fetal liver and adult bone marrow, approximately 1/3 of the neutrophils were early precursors (progenitors, preNeus and proNeus), and approximately 2/3 were immature neutrophils. In blood, a few precursors were found in E17.5 and NB mice, but not in adults, and the proportion of mature neutrophils increased progressively with age, culminating in the majority of neutrophils in adult blood being fully mature (Fig. 3E). In the embryonic and NB kidney and testis, both immature and mature neutrophils and, interestingly, also a few progenitor cells were observed (Fig. 3E). Overall, our scRNA-seq analyses revealed that the neutrophil development trajectory in utero is linear, neutrophils are present in steady-state non-hematopoietic tissues, and although those tissue-resident neutrophils are dominated by immature neutrophils, a few progenitor and early precursor cells are also identifiable.

### Tissue-specific and developmental-stage-dependent factors uniquely shape neutrophil transcriptomic profiles

We next evaluated whether neutrophil transcriptional profiles were similar across embryonic tissues or more aligned with blood neutrophil profiles. Our results showed that at E17.5, neutrophils in the kidney and testis typically expressed less than 50 tissue-specific DEGs (Fig. 3F and S4A and Data S1). Immature neutrophils in the bone marrow and liver displayed a higher number of DEGs (366 and 564, respectively), of which the vast majority were downregulated, compared to blood neutrophils (Fig. 3F and S4A and Data S1). Gene Ontology analysis revealed that both upregulated and downregulated genes in embryonic tissue-resident neutrophils were predominantly linked to general biological processes, such as the regulation of aerobic respiration and other cellular metabolic processes (Fig. 3G and H and S4B and C). By the NB stage, these gene expression differences were vastly diminished (Fig. S4D and Data S1). This aligned with our observation that no cluster-to-tissue distribution patterns emerged in our scRNA-seq analyses.

We observed several DEGs in mature neutrophils in the hematopoietic organs and blood when comparing E17.5 and adult cells. (Fig. 4A). Mature embryonic neutrophils highly expressed stefin family (type 1 cystatins) genes such as *Stfa1*, *Sfta2,* and *Stfa3* (Fig. 4B and C and Data S1), which encode cytoplasmic inhibitors of cysteine proteases and play a crucial role in the cellular antiviral response (Mihelič et al., 2006). Similarly, Olfactomedin 4 (*Olfm4*, OLFM-4), another protease inhibitor, was expressed consistently in embryonic neutrophils from the pro-neutrophil stage to mature neutrophils. In contrast, in adults, *Olfm4* expression was limited to a small, specific neutrophil subpopulation (Fig. 4D), as previously reported (Alder et al., 2019). To validate these findings at the protein level, we profiled neutrophils from different tissues by flow cytometry. We found that 84.2% (±4.8) of the Ly-6G^+^ circulating neutrophils in E17.5 embryos expressed OLFM-4, whereas only 6.7% (±2.3) of circulating neutrophils in adult blood were OLFM-4 positive (Fig. 4F and G). Similarly, 76.3% (±4.8) and 7.6% (±2.0) of neutrophils in embryonic liver and adult bone marrow, respectively, were OLFM-4 positive (Fig. 4F and G).

**Figure 4.**
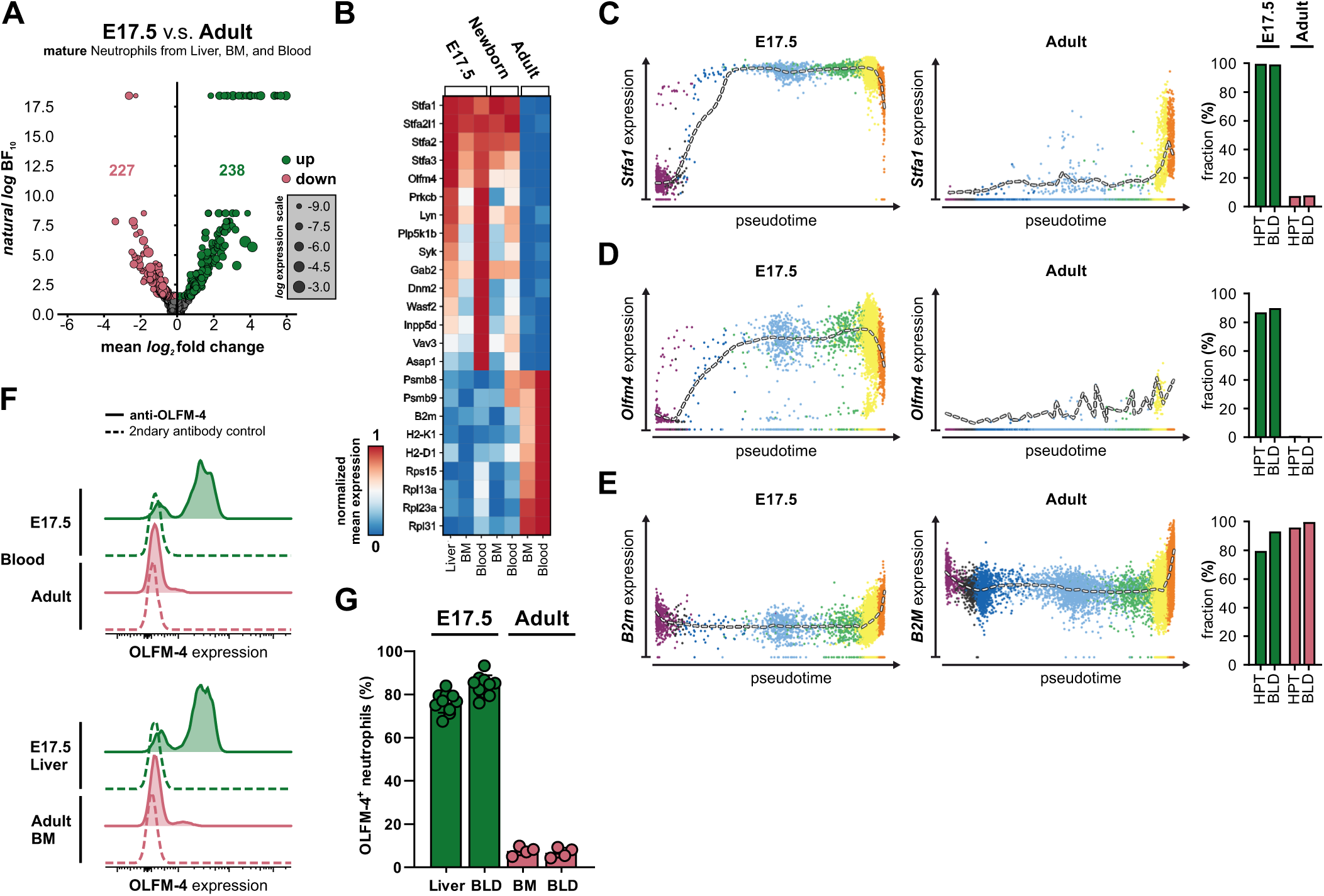
Embryonic and adult neutrophils have distinct gene expression signatures. **(A)** Volcano plot assessing differential gene expression between mature neutrophils from embryonic liver, bone marrow, and blood and adult bone marrow and blood. **(B)** Matrixplot of normalized expression for selected genes differentially expressed in mature neutrophils across ages and tissues. **(C-E)** Gene expression of *Stfa1*, *Olfm4*, and *B2m* along neutrophil maturation (pseudotime) and fraction of cells with non-zero gene counts (right) in hematopoietic tissues (HPT, fetal liver and bone marrow) and blood (BLD) in embryonic (left; green bars) and adult (right; red bars) neutrophils. **(F and G)** Representative histogram (F) and quantification (G) of OLFM-4 expression in embryonic (green) and adult (red) neutrophils. In G, each dot represents blood, liver, a pool of both embryonic femora and tibiae (BM) from one mouse or one adult femur from one mouse (n = 10 mice at E17.5 and n = 4 mice in adults). Data are presented as mean ± SD. Data from 3 independent experiments. BF, Bayes factor; BM, bone marrow.

Mature embryonic neutrophils in the blood highly expressed signaling molecules like *Lyn*, *Syk,* and *Vav3*, which were only weakly expressed in adult blood neutrophils (Fig. 4B). Genes down-regulated in embryonic neutrophils were often linked to antigen presentation on MHC-I, such as *B2m*, and cytoplasmic translation (Fig. 4B and E). We found that the genes upregulated in mature embryonic neutrophils often differed both by the fraction of positive cells and mean expression level, while the down-regulated ones only differed by the mean expression level. Moreover, the majority of DEGs observed in embryonic neutrophils were upregulated in newborn neutrophils when compared to adult cells (Fig. 4B and Data S1). We observed that immature neutrophils from E17.5 and adult mice also displayed multiple DEGs, albeit at a lower frequency than in mature cells. (Fig. 4B and S4E-G and Data S1). Collectively, our data show that embryonic and adult neutrophils express distinct gene programs and reveal new transcriptomic (*Stfas*) and protein (OLFM-4) markers selectively expressed in embryonic-type neutrophils.

### Embryonic neutrophils exhibit enhanced proliferation and metabolic flexibility

In adults, mature neutrophils are thought to be non-proliferative, short-lived, metabolically active cells that are mainly cleared by apoptosis (Aroca-Crevillén et al., 2024). We determined the proliferative capacity of neutrophils in embryos and adults by analyzing more closely the two-hour time point after BrdU pulse in various embryonic tissues (Fig. 5A and S3G). We found the biggest fraction of BrdU^+^ S-phase neutrophils in E17.5 liver, in which, on average, 10.3% (± 3.2) of immature neutrophils proliferated (Fig. 5A). This was almost two times more than in adult bone marrow (5.5% ± 0.9). In embryonic liver, but not in adult bone marrow, also mature BrdU^+^ neutrophils were observed. Also, embryonic testis and kidneys harbored a small number of proliferating immature neutrophils (Fig. 5A). To assess the viability of embryonic neutrophils, we conducted flow cytometry analyses of Annexin V staining and Fixable Viability Dye (FVD) binding (Fig. 5B). We found that 60-80% of neutrophils isolated from E17.5 liver, bone marrow, kidney, and testis were FVD^−^Annexin V^−^, which was comparable to the frequency of viable adult bone marrow neutrophils. However, more neutrophils in embryonic liver and kidney than in adult bone marrow were in a late apoptotic state (FVD^+^Annexin V^+^) (Fig. 5B).

**Figure 5.**
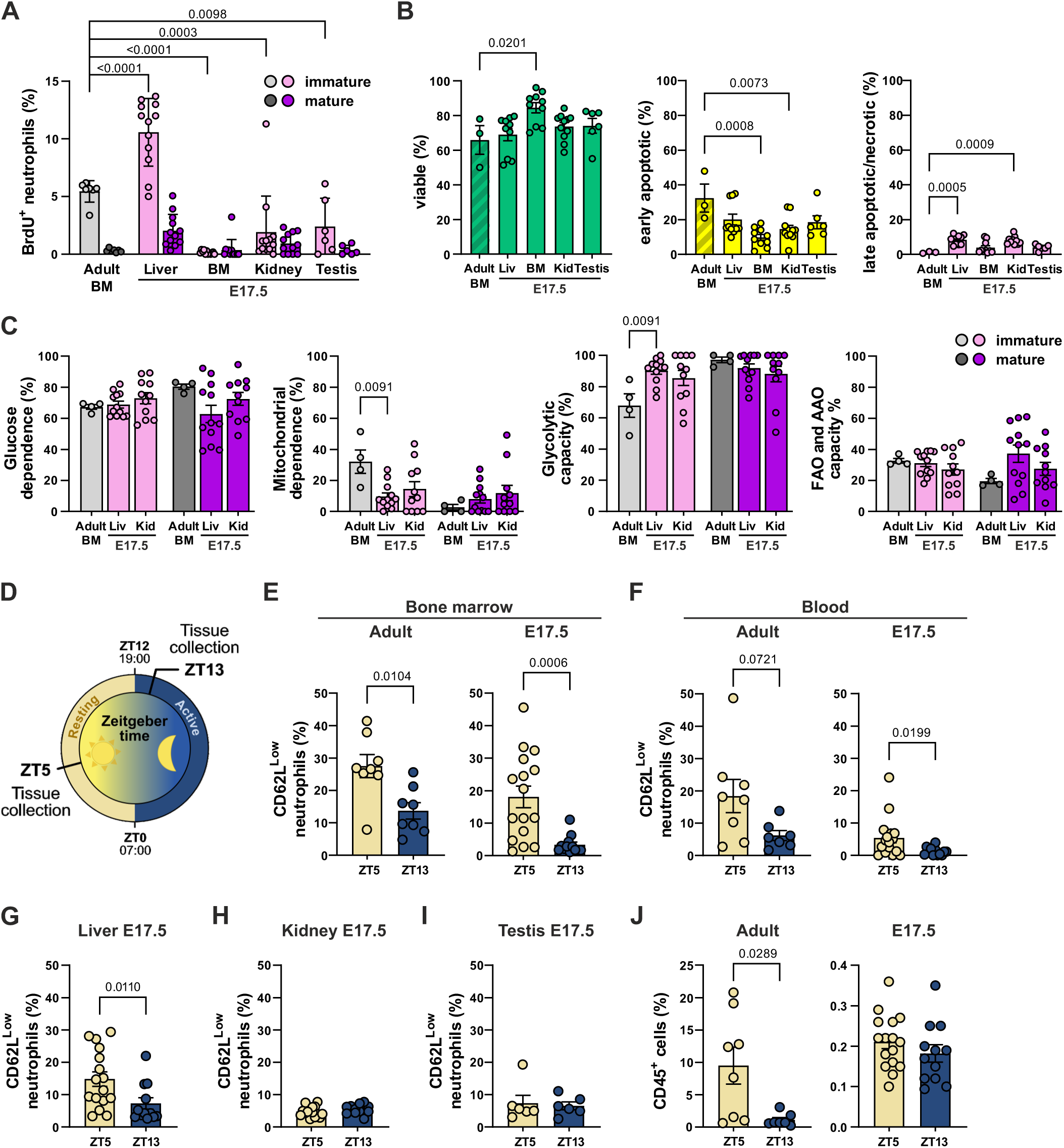
Embryonic neutrophils differ from adult neutrophils in proliferative and metabolic capacity and in diurnal rhythmicity. **(A)** Frequency of BrdU^+^ immature and mature neutrophils from adult bone marrow and indicated embryonic tissues in S-phase determined by BrdU uptake after 2 h circulation. **(B)** Frequency of viable (Annexin V^-^FVD^-^), early apoptotic (Annexin V^+^FVD^-^), and late apoptotic/necrotic (Annexin V^+^FVD^+^) neutrophils in adult bone marrow and in indicated embryonic (E17.5) tissues. **(C)** Metabolic profile of adult bone marrow and E17.5 liver (Liv) and kidney (Kid) -derived immature (light) and mature (dark) neutrophils. **(D)** Schematic overview of the circadian oscillation experiment. **(E-I)** Frequency of CD62L^low^ Ly-6G^+^ neutrophils in bone marrow (E) and blood (F) of adult and E17.5 tissues and in E17.5 liver (G), kidney (H), and testis (I) at Zeitgeber times 5 (ZT5) and 13 (ZT13). **(J)** Frequency of CD45^+^ leukocytes in adult and E17.5 blood at ZT5 and ZT13. In A-C and E-J, each dot represents one mouse, a pool of both femora and tibiae (BM) or a pool of both kidneys of one embryo, except for C, where in kidney, each dot represents a pool of 3 kidneys, and A, B, I, where in testis, each dot represents two males (4 testes). In A, mean ±SD are shown, and in B, C, E-J, mean ±SEM are shown. (A and B) two-way ANOVA with Dunnett’s multiple comparisons test, (C) two-way ANOVA with Tukey’s multiple comparisons test, and (E-J) Mann-Whitney test. All data are from 3-4 independent experiments. FVD, fixable viability dye; FAO, free acid oxidation; AAO, amino acid oxidation.

E17.5 tissue-resident neutrophils showed oxidative phosphorylation and mitochondrial ATP synthesis as the top upregulated gene pathways compared to blood neutrophils (Fig. 3G). To functionally assess the metabolic activity of embryonic neutrophils in the E17.5 liver and kidney, we took advantage of the flow cytometry-based single-cell method SCENITH (Argüello et al., 2020). We found that embryonic neutrophils relied primarily on glycolysis when glucose is readily available (Fig. 5C). No significant difference was observed between embryonic and adult neutrophils in their capacity to utilize fatty acids and amino acids as alternative energy sources when glucose oxidation was blocked (Fig. 5C). Consistent with previous findings (Riffelmacher et al., 2017), adult bone marrow-derived immature neutrophils exhibited a greater reliance on mitochondrial oxidative phosphorylation (OXPHOS) compared to mature neutrophils. However, this dependency was not observed in embryonic neutrophils (Fig. 5C). Notably, the embryonic immature cells had a better glycolytic capacity than adult immature cells to sustain protein synthesis when mitochondrial OXPHOS was inhibited (Fig. 5C). Collectively, our analyses indicate that embryonic neutrophils exhibit enhanced glycolytic capacity, different kinetics in progression of apoptosis compared to adult cells and a higher proliferative capacity. Moreover, proliferating tissue-resident neutrophils also existed in embryos.

### Distinct diurnal rhythms of embryonic neutrophils

In adults, the number of circulating neutrophils follows circadian oscillations (Adrover et al., 2019; Casanova-Acebes et al., 2013) The cells can be divided into CD62L^high^ “fresh” neutrophils, recently released from the bone marrow to blood stream, and CD62L^low^ “aged” neutrophils returning back to the bone marrow. Since the functionality of circadian cues in utero has been studied much less (Ono et al., 2021), we explored whether embryonic neutrophils exhibit circadian properties. Previous studies have shown that in adult blood, the highest frequency of aged neutrophils is found at Zeitgeber time 5 (ZT5, i.e., 5 h after turning lights on) and the lowest frequency at ZT13-ZT17 (Casanova-Acebes et al., 2013). Therefore, we collected tissues from E17.5 embryos at ZT5 and ZT13 for flow cytometric analyses (Fig. 5D).

We observed a fluctuation in the frequency of aged CD62L^low^ neutrophils according to the light-dark rhythm in hematopoietic tissues in embryos (Fig. 5E-G). A higher number of aged neutrophils were observed at the resting period (ZT5) compared to the beginning of the dark (i.e., active period) at ZT13 in bone marrow, blood, and liver (Fig. 5E-G). In contrast, the frequencies of aged neutrophils in the testis and kidney were unaffected by the circadian rhythm (Fig. 5H and I). Interestingly, the total leukocyte frequency (out of live cells) remained stable from ZT5 to ZT13 in embryonic blood, unlike in adults (Fig. 5J). These observations suggest that the circadian oscillation of neutrophil circulation and clearance are differentially regulated in hematopoietic and non-hematopoietic organs in utero.

### Embryonic neutrophils have the capacity for lineage-specific functions

To investigate lineage-specific functions of embryonic neutrophils, we conducted a set of in vitro experiments to measure phagocytosis, ROS production, and capability for neutrophil extracellular trap (NET) formation. To measure phagocytosis, we first incubated 0.5 µm fluorescent polystyrene beads with a single-cell suspension prepared from embryonic bone marrow, liver, and kidneys, and from adult bone marrow as a control (Fig. 6A). Flow cytometric analyses revealed that a lower percentage of embryonic bone marrow-derived mature neutrophils were loaded with beads compared to adult bone marrow derived cells. Moreover, mature fetal neutrophils isolated from liver, bone marrow, and kidney had comparable phagocytic capacity (Fig. 6B). Immature neutrophils in both embryos and adults were poorly phagocytic. (Fig. 6B).

**Figure 6.**
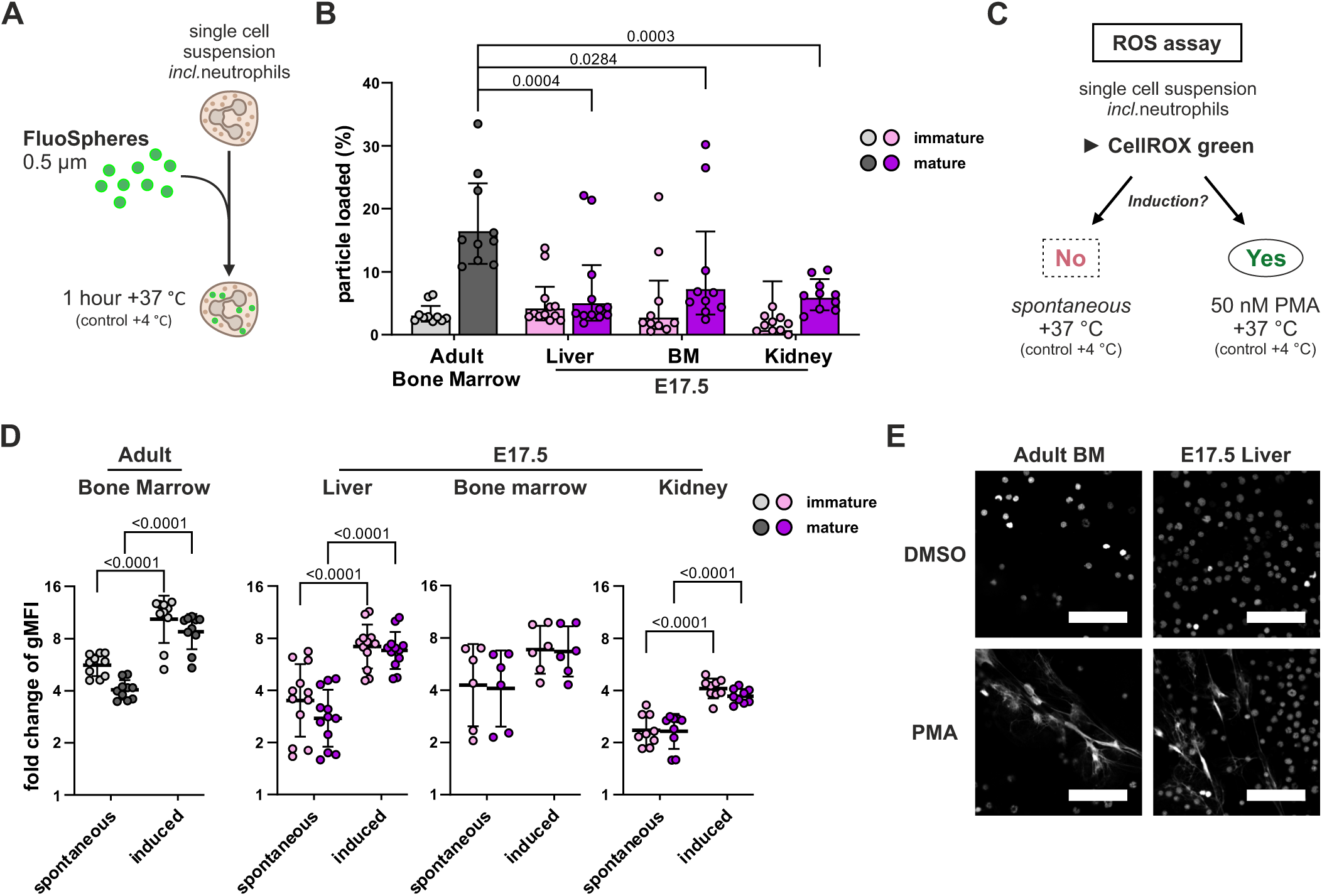
Embryonic neutrophils perform lineage-specific immune functions. **(A)** Schematic overview of the phagocytosis experiment. **(B)** Flow cytometric analyses of particle uptake by *in vitro* cultured neutrophils isolated from embryonic liver, bone marrow, and kidney, and adult bone marrow. Shown are frequencies of immature (light magenta, embryonic; light grey, adult) and mature (magenta, embryonic; dark grey, adult) neutrophils with ingested particle load at the indicated time points. **(C and D)** Analysis of ROS production in neutrophils. Experimental outline (C) and fold change of CellROX green signal in immature and mature neutrophils without (spontaneous) or with PMA induction (induced) (D) in cells isolated from embryonic liver, bone marrow (BM), and kidney (right) and adult BM (left) are shown. **(E)** Representative confocal immunofluorescence microscopy images of NET-like structures by leukocytes isolated from E17.5 liver (right) and adult BM (left) treated with either vehicle (DMSO; top) or induced with phorbol 12-myristate 13-acetate (PMA; bottom) and stained for DNA (gray). Scale bars 50 µm. In B and D, each dot represents one mouse (adult BM and embryonic liver), a pool of both femora and tibiae (BM), or a pool of both kidneys of one embryo. (B) Mixed-effects analysis with Tukey’s multiple comparisons test. (D) Two-way ANOVA with Šídák’s multiple comparisons test. In B and D, the geometric mean with ± geometric SD is shown. The data in B and D are from 3-4 independent experiments. BM, bone marrow; gMFI, geometric mean fluorescence intensity.

ROS production is one canonical outcome of phagolysosome activation in neutrophils (Wright et al., 2010). We therefore analyzed spontaneous and phorbol myristate acetate-induced (PMA) ROS production capacity in immature and mature embryonic neutrophils using the CellROX green-based flow cytometric assay (Fig. 6C). We found comparable basal ROS production in immature and mature neutrophils derived from embryonic bone marrow, liver, and kidney (Fig. 6D). Similar to adult bone marrow neutrophils, a significantly higher CellROX green signal was observed after PMA induction in immature and mature embryonic bone marrow, liver, and kidney neutrophils (Fig. 6D). Since we observed differences in apoptosis of embryonic and adult neutrophils, we also evaluated the potential of these cells to form NETs. Using PMA-stimulated release of DNA as a read-out, we observed comparable NET-like structures in cells from E17.5 embryonic liver and adult bone marrow (Fig. 6E). Collectively, these functional analyses show that embryonic neutrophils are endowed with effector functions like phagocytosis, ROS production, and NETosis both in hematopoietic and non-hematopoietic organs.

### Embryonic neutrophils respond to maternal immunosuppression and stimulation

Glucocorticoids are routinely used to enhance lung maturation in preterm infants (Bolt et al., 2001). Since glucocorticoids are also a powerful immunosuppressant, we investigated their effects on embryonic neutrophils. To detect the acute effect of DEX on embryonic neutrophils, we administered dexamethasone (DEX; 5 mg/kg) or PBS intraperitoneally (i.p.) to the pregnant dam at E17.5 (Fig. S5A) and we collected the blood and kidneys 2h after the DEX treatment for scRNA-seq analyses (Fig. S5A, B, and Data S2). DEGs analyses showed that in both tissues, immature and mature neutrophils exhibited reduced expression of genes associated with inflammatory processes. The down-regulated genes included those regulating neutrophil differentiation (*Lrrc25, Hist1h1e, Alas1*) (Junwei Lian et al., 2018; Liu et al.; Sollberger et al., 2020), recruitment (*Eno1, Orai1, Gadd45a*) (Sogkas et al., 2015; Salerno et al., 2012; Lu et al., 2025), and NET formation (*Il1b, Clec5a*) (Chen et al., 2017; Meher et al., 2018), whereas the expression of several genes with anti-inflammatory functions (*Arrb2, Gpr141, Cited2, Nfkbia, Dusp1, Il1r2, Tsc22d3*) (Espinasse et al., 2015; Tan et al., 2022; Pong Ng et al., 2020; Gaffal et al., 2014) was markedly elevated (Fig.S5D and E).

Both low-grade chronic inflammation (e.g., maternal obesity, gestational diabetes) and acute inflammatory conditions (e.g., bacterial infections) of the mother can affect the fetus (Pantham et al., 2015; Kumar et al., 2022). We used lipopolysaccharide (LPS)-induced gestational inflammation to study the inflammatory responses of embryonic neutrophils (Fig. S5F-H). To induce inflammation, pregnant dams were administered i.p. 100 µg/kg of LPS at E17.5, and tissues were collected 4 hours post-treatment for scRNA-seq analyses. While LPS exposure did not significantly alter neutrophil gene expression (Data S2), it resulted in a marked increase in the proportion of mature versus immature neutrophils (S5G and H) compared to both steady-state conditions (Fig. 3E) and DEX-treated mice (Fig. S5B and C). Overall, the results reveal that embryonic neutrophils are responsive to external stimuli, such as glucocorticoids and endotoxin, suggesting that embryonic neutrophils may serve as a reservoir ready to provide innate immune protection to the fetus in utero and at birth.

### Gfi1 genetic model reveals efficient neutrophil depletion compared to antibody and Cre-based strategies

To examine the effects of neutropenia in embryonic tissues, we initially applied anti-Ly-6G antibody (clone 1A8), which is commonly used in adults to specifically deplete neutrophils (Daley et al., 2008). We administered 1 mg of anti-Ly-6G antibody i.p. to pregnant females at E12.5, E14.5, and E16.5. After ex vivo extracellular Ly-6G staining using a fluorochrome-conjugated anti-Ly-6G antibody (clone 1A8), we found a significant absence of Ly-6G⁺ neutrophils in the liver of E17.5 embryos (Fig. S6A). However, when we used a secondary antibody ex vivo to detect the in vivo injected anti-Ly-6G antibody, we observed that Ly-6G^+^ neutrophils were not depleted but instead became undetectable by conventional direct Ly-6G surface staining due to antigen masking (Fig. S6A). Since Ly-6G depletion has been reported to be improved by “isotype switch” strategy, a dual antibody-based depletion approach (Boivin et al., 2020; Mukherjee et al., 2025), we further injected pregnant mice daily from E12.5 to E16.5 with anti-Ly-6G antibody (250 µg on the first day, followed by 100 µg on subsequent days). In parallel, an anti-rat IgG2a antibody (100 µg) was administered every other day starting at E12.5 (Fig. S6B). However, we still found numerous Ly-6G^+^ cells in the livers of the E17.5 embryos (Fig. S6B), which led us to conclude that the anti-Ly-6G antibody is an inefficient tool for neutrophil depletion in embryos.

Next, we crossed neutrophil-specific *Ly6g*^Cre-tdT/Cre-tdT^ mice with DTA^lox/lox^ (Ly6gCre-DTA) mice to enable the conditional expression of diphtheria toxin in neutrophils. However, only a ∼50% reduction in the frequency of Ly-6G^+^ blood leukocytes was observed in Ly6gCre-DTA mice, and the depletion efficacy in tissue neutrophils was even lower (Fig. S6C). This partial ablation may be attributed to the presence of Ly-6G^low^ neutrophil or Ly-6G^negative^ pre-neutrophil populations in the tissues that escape Cre-mediated recombination because of the low efficiency of *Ly6g*^+/Cre-tdT^ mice (Fig. S2B).

To achieve more extensive neutrophil depletion, we took advantage of the previously generated *Gfi1^R412X/R412X^* (hereafter called *Gfi1^mut/mut^*) mouse model (Muench et al., 2018). Growth Factor Independence 1 (*Gfi1*) is a transcription factor regulating the differentiation of neutrophils from their progenitors (Vassen et al., 2007; Van Der Meer et al., 2010), directing the pre-T-cell differentiation (Yücel et al., 2004; Fraszczak and Möröy, 2021), formation of inner ear hair cells and pancreatic acinar/centroacinar unit (Fiolka et al., 2006; Wallis et al., 2003; Qu et al., 2015), and the development of secretory cell types in the intestine (Shroyer et al., 2005).

While complete knockout of *Gfi-1* affects multiple leukocyte lineages and disrupts development in various cell types, the *Gfi1^R412X^* mutation, which introduces a premature stop codon, selectively only prevents the formation of mature neutrophils in postnatal mice (Muench et al., 2018, 2020a). We observed that 4-week-old *Gfi1^mut/mut^* mice showed reduction of bone marrow and blood Ly-6G^+^ neutrophils (6.6% and 3.7%, respectively; compared to 21.4% and 8.6%, respectively, in wild-type controls (Fig. S6D)) and monocytosis (Fig. S6E). We discovered that at E17.5, *Gfi1^mut/mut^* embryos exhibited a more pronounced reduction in neutrophil numbers compared to adult mutants, with a significant decrease observed both in hematopoietic and peripheral tissues (Fig. 7A, S6F, and S6G). A mild monocytosis was also detected in the E17.5 peripheral blood (Fig. S6H). Of note, in embryos, but not in adults (Fig. S6D), the neutrophil levels were reduced to a smaller extent, but without any concomitant alterations in monocytes, in heterozygous *Gfi1^+/mut^*mice (Fig. S6F-H).

**Figure 7.**
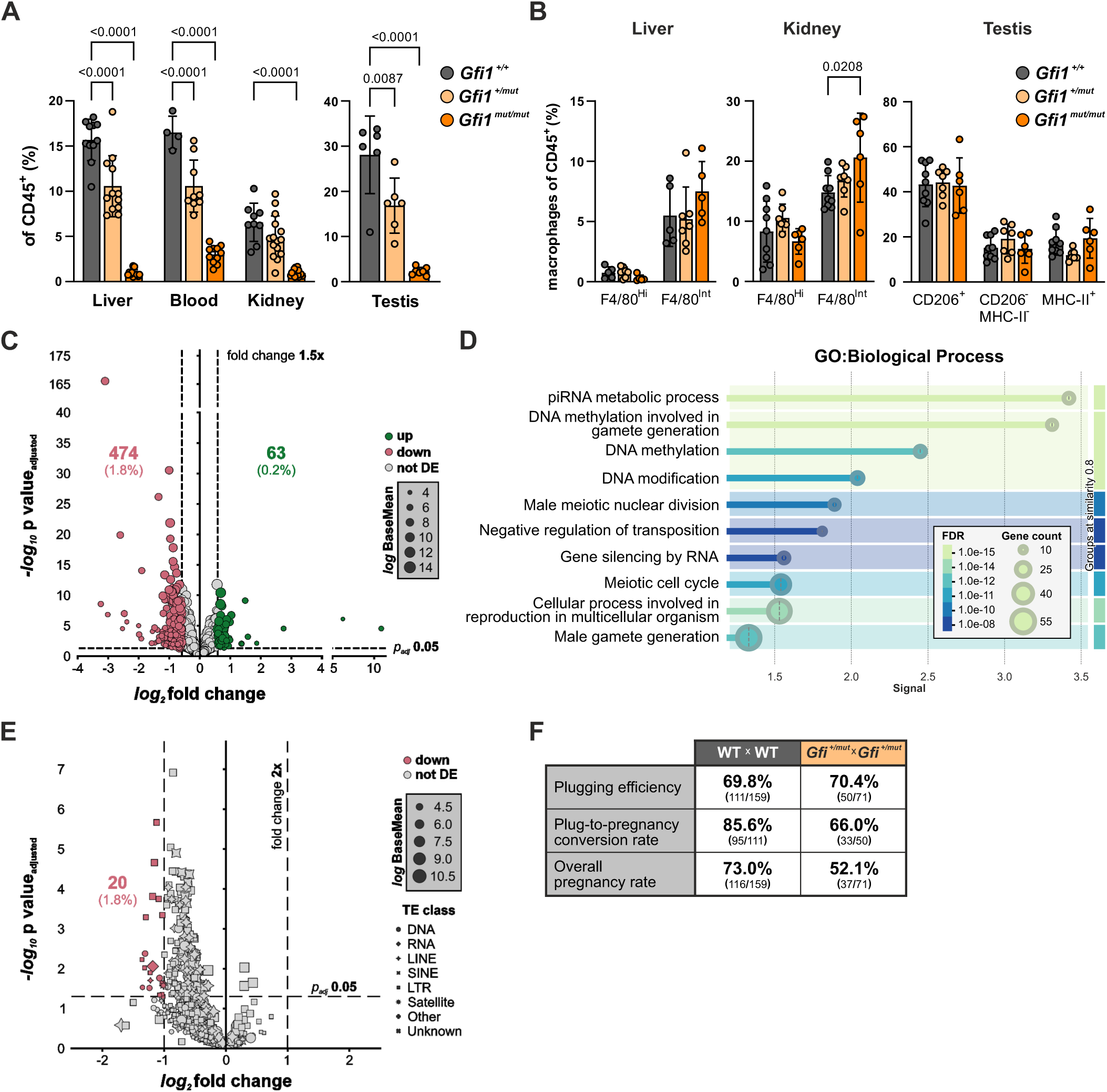
Genetic neutrophil deficiency results in altered testicular transcriptome and reproduction. **(A)** Flow cytometric analyses of neutrophils in the indicated embryonic (E17.5) tissues of *Gfi1^+/+^* (gray), *Gfi1^+/mut^* (light orange), and *Gfi1^mut/mut^* (orange) mice. **(B)** Flow cytometric analyses of resident macrophage populations in the indicated tissues of adult *Gfi1^+/+^*(gray), *Gfi1^+/mut^*(light orange), and *Gfi1^mut/mut^* (orange) mice. **(C)** Volcano plot assessing differential gene expression in testes of *Gfi1^mut/mut^* and WT embryos at E17.5 as determined by bulk RNA-sequencing. **(D)** STRING functional enrichment visualization of the top 10 gene ontology terms related to biological processes enriched in downregulated DEGs in the testes of *Gfi1^mut/mut^* embryos at E17.5. **(E)** Volcano plot of differential expression of transposable elements (TE) in testes between *Gfi1^mut/mut^*and WT embryos at E17.5 as determined by bulk RNA-sequencing. **(F)** Analysis of breeding characteristics of *Gfi1^+/mut^* x *Gfi1^+/mut^* crossings compared to crossings of WT animals. Shown are percentage and absolute numbers (parentheses) of observed overall rate of pregnancies, vaginal plugging efficiency, and rate of pregnancies after detection of a vaginal plug. In A, each dot represents liver, blood, a pool of both kidneys, or a pool of both testes from one mouse. In B, each dot represents the liver, one kidney, or one testis from one mouse. Data in A and B are presented as mean ± SD and are from 3-5 independent experiments. Statistical Analysis was performed using Two-way ANOVA with Dunnett’s multiple comparisons test (A and B). FDR, false discover rate corrected p-values, signal, weighted harmonic mean between the observed/expected ratio and -log(FDR), LINE, long interspersed nuclear element; SINE, short interspersed nuclear element; LTR, long terminal repeat.

### Neutrophil depletion disturbs the expression of the piRNA pathway in the testis

During immune cell differentiation, chromatin undergoes extensive reprogramming. In adults, neutrophil serine proteases (NSPs), including cathepsin G (CTSG), neutrophil elastase (ELANE), and proteinase 3 (PRTN3), have been shown to regulate histone modifications during monocyte-to-macrophage differentiation (Cheung et al., 2021). Many tissue-resident macrophage populations are established before birth, primarily from fetal liver-derived monocytes (Hoeffel et al., 2015; Rantakari et al., 2016; Guilliams et al., 2013; Sheng et al., 2015). To assess whether the absence of neutrophils in *Gfi1^mut/mut^* embryos affects the establishment of tissue-resident macrophage populations, we performed flow cytometry analysis on 4-week-old *Gfi1^mut/mut^* mice, focusing on the organs known to harbor macrophages derived from embryonic liver monocytes (Lokka et al., 2020; Schulz et al., 2012; Hoeffel et al., 2015). However, no difference in macrophage frequencies was observed between adult *Gfi1^mut/mut^* and control mice in liver, kidney, or testis, apart from a minor increase of F4/80^intermediate^ macrophages in the kidney of *Gfi1^mut/mut^* mice (Fig. 7B).

To further evaluate the impact of neutrophil reduction on general tissue development and function, we performed bulk RNA sequencing on the E17.5 testes and kidneys of *Gfi1^mut/mut^* and control mice. The kidney showed only minor changes in gene expression between the two genotypes (Fig. S6I and Data S3). In contrast, we found 63 upregulated and 474 downregulated genes in the testis of *Gfi1^mut/mut^* (Fig. 7C and Data S3). Among upregulated genes, gene enrichment analysis indicated cholesterol metabolism and steroid biosynthesis as the major hits in the *Gfi1^mut/mut^* (Fig. S6J). Interestingly, genes downregulated in the embryonic *Gfi1^mut/mut^*testis included many associated with the metabolic processing of P-element-induced wimpy testis (PIWI)-interacting RNAs (e.g., *Tdrd12*, *Asz1*, *Gpat2*, and *Fkbp6*) and DNA methylation (e.g., *Morc1*) (Data S3), both critical for testicular development and spermatogenesis in mammals (Kuramochi-Miyagawa et al., 2008; Luteijn and Ketting, 2013). Gene enrichment analyses also confirmed the downregulation of the piRNA metabolic process and DNA methylation in *Gfi1^mut/mut^*testes (Fig. 7D and S6K).

In the testis, piRNAs are predominantly expressed in germline cells. During fetal development, their expression coincides with the period of germline DNA methylation reprogramming, where they play a crucial role in the transcriptional and post-transcriptional silencing of retrotransposons (Kuramochi-Miyagawa et al., 2008; Reznik et al., 2019). Defective function of the piRNA pathway can lead to the activation of retrotransposon expression, including the long interspersed nuclear elements (LINEs). However, we saw no major difference in LINE1 expression in *Gfi1^mut/mut^*E17.5 testis compared to control (Fig.S6L). In broader analyses of transposable elements in the *Gfi1^mut/mut^* E17.5 testis, most piRNAs targeting LINEs, Short interspersed nuclear elements (SINEs), and Long terminal repeat (LTR) elements were not upregulated, but instead were downregulated (Fig. 7E and Data S3).

Gonadal piRNAs are vital to normal spermatogenesis, and disruption of the piRNA pathway machinery has been linked to male infertility even in humans (Kuramochi-Miyagawa et al., 2008; Luteijn and Ketting, 2013; Stallmeyer et al., 2024). Histological examination of *Gfi1^mut/mut^* testes at E17.5 and 7 weeks of age did not reveal apparent morphological abnormalities (Fig. S6M) and the occurrence of vaginal plugs was similar between *Gfi1^+/mut^* and control females in fertility assays (Fig.7F). Nevertheless, the rate of successful pregnancies following plug detection (WT 73.0% *vs Gfi1^+/mut^* 52.1%) and the overall pregnancy rate (WT 85.6% *vs Gfi1^+/mut^* 66.0%) was compromised in the *Gfi1^+/mut^* group compared to controls (Fig.7F). In conclusion, these findings imply that in neutropenic *Gf1^mut/mut^* mice the number of tissue-resident neutrophils in embryonic testis is reduced and this is associated with defective regulation of piRNA and methylation programs and subnormal fertility.

## Discussion

Neutrophil biology in non-hematopoietic organs under steady-state conditions and especially in fetuses has remained virtually unexplored. Here, we demonstrate kinetically shifting neutrophil infiltrates in multiple non-hematopoietic tissues during embryogenesis under homeostatic conditions. Notably, fetal neutrophils predominantly resided within the parenchyma of tissue rather than within the vasculature. The tissue-resident embryonic neutrophils were transcriptionally different from circulating embryonic neutrophils and from adult neutrophils. We also showed that the embryonic neutrophils responded to external immunostimulatory and immunosuppressive cues at transcriptional and functional levels. In addition to their innate immunity roles, analyses of neutrophil-deficient mice suggested that embryonic tissue-resident neutrophils may additionally display homeostatic functions, such as regulation of fertility.

In adults, neutrophils exhibit pronounced circadian rhythmicity, with tissue-resident neutrophils increasing in number as their presence in circulation declines (Ella et al., 2016; He et al., 2018). We found a higher proportion of aged neutrophils during the light phase in hematopoietic tissues of embryos. This is consistent with observations showing that mammalian embryos follow the circadian clock before birth, primarily due to maternal cues, such as varying levels of melatonin, dopamine, and glucocorticoids (Ono et al., 2021). Interestingly, this oscillatory pattern was not found in non-hematopoietic tissues such as the testis and kidney, where the frequencies of aged neutrophils remained constant throughout the circadian cycle. In embryos, the tissue-resident neutrophils displayed multiple DEGs compared to blood-borne neutrophils. Furthermore, our intravascular administration of fluorescently labeled Ly-6G antibody into the umbilical veins and the whole-mount imaging revealed that many tissue-resident neutrophils in embryos were at extravascular parenchymal locations rather than intravascularly under steady-state conditions. Collectively, our data suggests that embryonic tissue-resident neutrophils in non-hematopoietic organs at steady-state should be considered as a separate cell type distinct from circulating blood neutrophils.

Transcriptomic analysis showed that embryonic neutrophils follow a linear developmental pathway from progenitor cells via pro- and pre-neutrophils and immature neutrophils into mature neutrophils. In general, fetal tissues harbored a higher proportion of immature neutrophils, many of which were actively proliferating, compared to adults. In adults, low numbers of bone marrow-derived neutrophils infiltrate almost every naive tissue except the reproductive organs and the brain (Puga et al., 2012; Casanova-Acebes et al., 2013). Interestingly, in the embryonic testis, which is formed from the aorta-gonad-mesonephros structure during early hematopoiesis, we identified a few tissue-resident neutrophil progenitor cells. This finding raises the possibility that some of the tissue-resident neutrophils in the testis may be derived from local neutropoiesis rather than from extravasated blood-borne neutrophils. Otherwise, it would imply that circulating neutrophil progenitors and immature neutrophils would have effective machinery for executing the multistep leukocyte extravasation cascade even in a non-inflamed steady-state environment, in which vascular endothelial cells usually lack the majority of adhesion and attractant molecules.

Tissue-resident neutrophils displayed features compatible with dormant immune functions. For instance, the translation of MHCI and other molecules involved in antigen processing was reduced compared to adult neutrophils. Moreover, members of the Stefin family were highly abundant in fetal neutrophils, whereas they were practically absent from adult neutrophils. Stefins are poorly characterized proteins known as intracellular cysteine protease inhibitors (Mihelič et al., 2006), suggesting pronounced control of lytic activities in embryonic neutrophil cells. We also showed both at the mRNA and protein level that OLFM4 expression is characteristic of embryonic but not adult neutrophils. In other cell types, OLFM4 has been reported to be anti-apoptotic and to induce the expression of PD-L1 (Liu et al., 2018, 2012; Lin et al., 2022), implying that its expression in embryonic neutrophils would also be compatible with reduced immunoreactivity.

Although short-lived, neutrophils in adults can polarize to different types, as exemplified by the identification of anti-inflammatory N2-type neutrophils in the tumor microenvironment (Fridlender et al., 2009; Andzinski et al., 2016; Casbon et al., 2015). Among macrophages, fetal-derived tissue-resident macrophages share several common features with antitumor or M2-like macrophages (Park et al., 2022). Therefore, it will be interesting to study in the future if immunosuppressive N2-neutrophils in adults share common features with embryonic neutrophils. Nevertheless, our functional phagocytosis, ROS, and NET production assays showed that embryonic neutrophils have an intact machinery for executing effector functions in response to activation signals. For instance, embryonic neutrophils showed prompt responses to bacterial immunostimulants (LPS), which readily pass from the mother to the fetus through the placenta (Brown et al., 2019; Simões et al., 2018). These observations suggest that embryonic neutrophils may be tuned towards immunosuppressive reaction pathways, but that they are fully competent to respond to potential microbial and other challenges during intrauterine development and at birth.

The possible homeostatic roles of neutrophils in adults have remained largely enigmatic. However, recent research has revealed that neutrophils can also perform important homeostatic functions. In the skin, for example, matrix-producing neutrophils contribute to the composition and structure of the extracellular matrix, promoting the barrier function of the skin (Vicanolo et al., 2025). Using the genetic *Gfi1^mut/mut^* mouse model, we observed significant differences in the transcriptome of the embryonic testis in the presence and absence of neutrophils. More specifically, gene pathways related to steroid synthesis and piRNA regulation were significantly altered. Importantly, macrophage numbers in testes were not affected in *Gfi1^mut/mut^* mice. These data suggest that neutrophils can also support homeostatic, non-immunological functions under steady-state conditions.

In conclusion, our data provides a single-cell atlas of neutrophils in hematopoietic and non-hematopoietic organs under steady-state and perturbed conditions in embryonic and newborn mice. Functionally, we show the presence of extravascular neutrophils in normal tissues and transcriptomic differences between blood-borne and tissue-resident neutrophils. We demonstrate that embryonic neutrophils are responsive to danger stimuli and capable of executing several prototypical neutrophil immune functions. Neutrophils in non-hematopoietic tissues also contributed to homeostatic, non-immunological functions in growing mice, suggesting a broader role than expected for this leukocyte type under normal and pathological conditions.

## Materials and methods

### Mice

C57BL/6N WT mice were acquired from Janvier Labs. The C57BL/6-*Ly6g^tm2621^ ^(Cre-tdTomato)^ ^Arte^* reporter mice were kindly provided by Prof. Matthias Gunzer (Hasenberg et al., 2015). These mice were crossed with B6.Cg-*Gt(ROSA)26Sor^tm14(CAG-tdTomato)Hze^*/J for enhanced tdTomato expression. B6.129P2-Gt(ROSA)26Sortm1(DTA)Lky/J diphtheria toxin-producing mice were kindly provided by Prof. Pipsa Saharinen. Neutropenic *Gfi1^R412X/R412X^* mice (Muench et al., 2020) were acquired from Jax lab (*Gfi1^em1Hlg^*/J, 035156). All mice were kept in specific pathogen-free conditions under controlled environmental conditions (temperature 21±3 °C, 12 h light cycle) at the Central Animal Laboratory of the University of Turku (Turku, Finland). Mice had access to food and water *ad libitum*. In timed matings, the day when the vaginal plug was observed was recorded as embryonic day 0.5 (E0.5). Animal experiments were conducted under the revision and approval of the Regional Animal Experiment Board in Finland (license numbers 14685/2020 and 23546/2023) according to the 3R principles and the Finnish Act on Animal Experimentation (497/2013).

### Cell isolation

According to the recommendations of the Animal Experiment Board, adult mice were euthanized with carbon dioxide asphyxiation followed by cervical dislocation, newborn mice by decapitation, and one-week and two-week-old mice were terminally anesthetized with isoflurane before decapitation. Embryos were dissected from the uterus and euthanized by hypothermia followed by decapitation. Tissues were harvested around 8:00-10:00, except for the circadian rhythm studies, in which tissues were collected at Zeitgeber time 5 (ZT5, 12:00) and at ZT13 (20:00).

Blood was collected by cardiac puncture from euthanized adult mice, and by bleeding onto heparin-Phosphate Buffered Saline (PBS; D8537, Sigma-Aldrich; 50 µl of 100 IU/ml heparin (585661, LEO Pharma) in 500 µl of PBS) from euthanized E17.5 – 2-week-old mice. Erythrocytes were lysed from blood as described (Rantakari et al., 2015). Testes, kidneys, liver, thymus, lungs, pancreas, brain and spleen were gently minced with scissors and digested with 50 µg/ml DNase 1 (10104159001, Roche) and 1 mg/ml Collagenase D (11088866001, Roche) in Hank’s Buffered Saline (HBSS; RNBF2378, Sigma-Aldrich) solution at 37 °C for 15 to 45 min. After digestion, the brain cells were resuspended in isotonic Percoll (17-0891-01, GE Healthcare), and the microglia were isolated as described (Ginhoux et al., 2010). The kidneys and liver from 5- and 8-week-old mice were homogenized in RPMI-1640 (R5886, Sigma-Aldrich) medium in a gentleMACS C-tube (130-093-237, Miltenyi Biotech) with a Miltenyi Biotech gentleMACS Dissociator, and leukocytes were isolated by OptiprepTM (D1556, Sigma-Aldrich) gradient centrifugation. Bone marrow cells were collected by carefully crushing and rinsing the femora (and from embryos also tibiae) with HBSS, followed by sequential filtering through silk cloth (pore size 77 µm, 77-48Y115, Marabu Scandinavia) until no visible aggregates were detected. Finally, all the isolated cells were washed with 500 µl of PBS and suspended in PBS and filtered through a silk cloth before cytospin or labeling for cell sorting or flow cytometry.

### Flow cytometry

Prior to antibody staining, cells were stained with Fixable Viability Dye eFluor® 780 or 506 (65-0865-14 or 65-0866-14, respectively, Thermo Fisher) and Fc-receptor binding was blocked with anti-CD16/CD32 antibody (clone 2.4G2, BE0307, Bio X Cell). The cells were then incubated with fluorophore-conjugated primary antibodies diluted in flow cytometry buffer (PBS with 2% (v/v) fetal calf serum (FCS, S181B-500, Biowest) and 0.05% (v/v) sodium azide (S2002, Sigma-Aldrich)) at 4 °C for 30 min. Alternatively, unconjugated primary antibodies followed by appropriate fluorophore-conjugated secondary antibodies were used. To label intracellular antigens, either Foxp3 Transcription Factor Staining Buffer Set (00-5523-00, Thermo Fisher) or BrdU Flow Kit (559619, BD Pharmingen) was used according to the manufacturer’s protocol. After staining, the samples were fixed with 1% (v/v) formaldehyde (1040021000, Merck) in PBS unless otherwise indicated. In apoptosis experiments, the cells were washed with Annexin V binding buffer (422201, BioLegend) and incubated with BV421-conjugated Annexin V (640924, BioLegend) for 10 min to label apoptotic cells before fixation. All antibodies used in this study are listed in Data S5. Samples were acquired on a BD Biosciences LSRFortessa flow cytometer, and data were analyzed with FlowJo software (BD Biosciences).

### Single-cell cytology

Tissues were harvested and processed into a single cell suspension as described above. CD45^+^ Ly-6G^+^CD101-positive and -negative cells were sorted from the single-cell suspensions using a Sony SH800 Cell Sorter. The purified cells were cytospinned (Thermo Shandon Cytospin 3 Centrifuge) onto SuperFrost glass slides, air-fixed, and stained with modified Wright-Giemsa staining (9990701, Thermo Scientific). The slides were imaged with Olympus BX60 fluorescence microscope equipped with Olympus U-Plan Apochromat 100x/1.35 objective using Cargille Type B immersion oil. White-balance adjustments were done using Fiji/ImageJ (Schindelin et al., 2012).

### Wholemount tissue clearing and volumetric imaging

Sample preparation and tissue clearing were performed as described previously (Hofmann et al., 2021). Briefly, intact tissues were harvested from embryos at E17.5, initially fixed for 1 hour at room temperature in IC Fixation Buffer (00-8222-49, Thermo Fisher) followed by incubation in 25% (v/v) fixative in PBS overnight at 4 °C. Subsequently, tissues were blocked by incubation in Blocking buffer (PBS with 0.3% (v/v) Triton X-100 (T8787, Sigma-Aldrich), and 1% (v/v) each of FCS, bovine serum albumin (BSA; P6154, Biowest), and other normal sera according to the secondary antibody species (all Jackson ImmunoResearch)) overnight and stained with an appropriate mix of primary antibodies diluted in Blocking buffer for 72 to 96 hours. Samples requiring secondary antibody staining were rinsed with PBS, washed overnight in Blocking buffer, and stained with the appropriate secondary antibodies diluted in Blocking buffer for 32 to 36 hours. All antibodies used in this study are listed in Data S5. Finally, samples were washed (PBS with 0.2% (v/v) Triton X-100, and 0.5% (v/v) 1-thioglycerol (M6145, Sigma-Aldrich)) and then dehydrated using ascending dilutions of 30%, 50%, 70%, 100%, and 100% (v/v) isopropanol (I9516, Sigma-Aldrich) for 30 to 60 minutes each, and cleared using undiluted ethyl cinnamate (W243000, Sigma-Aldrich).

Imaging (for Figs. 2A, D-F and S2C-F) was performed with a 3i CSU-W1 spinning disk confocal microscope (50-µm pinholes) equipped with 405/488/561/610 nm solid state lasers combined with an illumination field uniformizer (except Fig. S2C), corresponding band-pass filters, a Teledyne Photometrics Prime BSI or Hamamatsu Orca-Flash4.0 sCMOS camera, Zeiss Plan-Apochromat 10x/0.45 Ph1, Plan-Apochromat 20x/0.8, or LD Plan-Neofluar 40x/0.6 Corr Ph2 objectives, and using 3i SlideBook microscope control software (ver. 6.0+). Alternatively, images (for Figs. 2B, C, and G and S2G and H) were acquired using a Leica STELLARIS 8 FALCON FLIM confocal laser scanning microscope (CLSM) equipped with a tunable white light laser and opto-acoustic filtering, Leica HC FLUOTAR L VISIR 16x/0.60 IMM incl BABB or HC PL APO 20x/0.75 IMM CORR CS2 objectives using Leica F Type immersion oil, and Power HyD X, HyD S, and HyD R detectors controlled by Leica LAS X software.

CLSM imaging was performed by 600 Hz bidirectional scanning of 2048x2048 points with 4x line averaging, 0.75-1x zoom at 1 A.U. (at 580 nm) pinhole size, system optimized z-step size, and frame sequential scans. CSLM images were processed using Leica LIGHTNING deconvolution (RIMedium=1.558, strategy=adaptive). In some images (Figs. 2C and G and S2H), minor antibody-mediated image artifacts in tdTomato staining were masked to improve image clarity using surface objects (smooth = false, intensity threshold = 2780.88 / 8000, voxels < 500 / 300 [Fig. 2C / 2G and S2H]) using Oxford Instruments IMARIS (ver. 10.0.2). All images were linearly adjusted for brightness and contrast and rendered in Oxford Instruments IMARIS Viewer (ver. 10.0+).

### Blood neutrophil labeling (umbilical cord injections)

To ensure the embryos are alive during antibody injection, the uterus of the pregnant dam was collected at E17.5 into pre-warmed PBS. Embryos were dissected onto individual 3 ml dishes containing warm PBS, and amniotic sacs were opened. Each embryo was injected in the umbilical vein with a total volume of 10 µl of antibody-PBS-Evans blue (containing 0.5 µg of Ly-6G-A647 antibody (127610, BioLegend) and 0.25% (w/v) Evans blue (E2129, Sigma-Aldrich)). The solution was allowed to circulate in the body for around one minute. After that, the embryo was euthanized, and blood was collected by bleeding into heparin-PBS. Kidneys, liver, testes, and femora and tibiae were collected and processed as described in Cell isolation.

### Single-cell RNA-sequencing

Single-cell suspensions were generated as described above from livers (only E17.5), bone marrow, blood, kidneys (both), testes (both) from WT E17.5 embryos (pooled samples: liver n = 30, bone marrow n = 4, blood n = 12, kidney n = 9, and testis n = 9) and NB animals (pooled samples: bone marrow n = 10, blood n = 9, kidney n = 9, and testis n = 9) in steady-state and after maternal dexamethasone treatment (pooled samples: blood n = 3 and kidney n = 3) or LPS exposure (pooled samples: blood n = 3 and kidney n = 5), [n, number of animals]. E17.5 liver and NB bone marrow samples were collected from both male and female embryos/animals. All other samples were collected only from male embryos/animals. Tissues for “newborn testis” sample originate from animals between E18.5 (c-sectioned) and postnatal day 1.

Cells were stained for flow cytometry without fixation as described above. The cells were sorted with a Sony SH800 Cell Sorter (100-µm nozzle, purity mode). Single cells were gated with FSC-H vs. FSC-W and SSC-H vs. SSC-W to avoid doublets, and dead cells were excluded. Live single CD45^+^ cells (kidney, testis) or CD45^+^ CD3^-^ CD45R^-^ cells (liver, BM, blood) were sorted into the RPMI-1640 medium containing 2% (v/v) FCS. Freshly sorted CD45^+^ cells were processed immediately according to 10X Genomics guidelines (CG000126_Guidelines for Optimal Sample Prep Flow Chart RevA). Single-cell RNA-sequencing libraries were prepared according to the manufacturer’s instructions (CG00052 RevB) using 10X Genomics Chromium Single Cell 30 Library and Gel Bead Kit v3 and Chromium Single Cell A Chip Kit. The libraries were sequenced using the Illumina Novaseq6000 Sequencing System (RRID: SCR_016387). The *Cell Ranger* (ver. 7.1.0 or 7.0.1, 10X Genomics) “count” function was used to align samples to the mm10-2020-A reference genome, quantify reads, and filter reads and barcodes. Data analysis and displays were generated using the *scanpy* (ver. 1.9.8, SCR_01813999), *scvi-tools* (ver. 1.0.4, SCR_026673), and other common Python (ver. 3.10+, SCR_008394) toolboxes.

The count matrices of our newly generated datasets were concatenated with selected, publicly archived, steady-state datasets generated using 10X Genomics technology of adult bone marrow and blood from Ballesteros et al. (Ballesteros et al., 2020) (GSE142754; “blood_ZT5”, “BM1”, “BM2”), *Tabula Muris* (Schaum et al., 2018) (GSE109774; “Marrow-10X_P7_2”, “Marrow-10X_P7_3”), and Xie et al. (Xie et al., 2020) (GSE137539; “wt_ctl_bm1”, “wt_ctl_bm2”, “wt_ctl_pb2”) (Fig. S3A).

Low-quality observations, filtered individually per sample based on observed distribution by number of expressed unique genes (minimum: 400–800, maximum: 2500–7000), total read counts (maximum: 10000-52500), and percentage of mitochondrial genes (maximum.: 12%–20%) and genes expressed by fewer than three cells were removed (Data S4). Counts per cell were normalized as total counts over all genes (target sum: 10^4) and were *log1p* transformed. Next, principal components (PCA) were calculated and a single cell Variational Inference (scVI) (Lopez et al., 2018) model (layer: “counts”, categorical covariates: “source”, “batch”, n_latent: 30, n_layers: 2) was generated. For the complete dataset, the neighborhood graph (nearest neighbors: 50) based on the scVI model’s latent space representation, Uniform Manifold Approximation and Projection (UMAP; SCR_018217103; min. distance: 0.5, neg. edge sample rate: 15), and initial *Leiden* clustering (ver. 0.10.1; resolution: 1.0) were computed with the indicated settings and initial cluster annotation was performed.

Under-resolved clusters containing hematopoietic and neutrophil-committed progenitors were selected and processed again using Leiden clustering (resolution: 0.75), resulting in the final overall cluster annotation (Fig. S3C). Markov affinity-based graph imputation of cells (van Dijk et al., 2018) as implemented in the *Palantir* (Setty et al., 2019) package (ver. 1.3.2) was used for improved visualization of gene expression, where indicated. Pseudotime analysis was performed with *scFates* (Faure et al., 2023) (ver. 1.0.6) by generating a principal tree (method: “ppt”, ppt_metric: “cosine”, Nodes: 300, ppt_sigma: 0.1, ppt_lambda: 50) and computing pseudotime values (n_map: 600) based on the distance of the projected cell position on the tree from the root node.

Differential expression genes were computed with *sci-tools*’s “differential_expression” function (mode: “change”, delta: 0.25, fdr_ target: 0.05) (Gayoso et al., 2022) and restricted to genes either expressed in at least 20% of cells in both or in more than 50% of cells in one of the comparison groups. Functional pathway enrichment analyses and visualizations were performed using the STRING database and its “Multiple Proteins by Names / Identifiers” search tool (ver. 12.0) (Szklarczyk et al., 2023).

### Cell proliferation assay

Pregnant dams at E17.5 were injected i.p. with 120 µl of 10 mg/ml 5-bromo-2’-deoxyuridine (BrdU, 51-2420KC, BD Biosciences) and euthanized 2, 24, 48, or 72 hours later. Tissues were processed and stained for flow cytometry with additional intracellular staining with BrdU Flow Kit according to the manufacturer’s protocol. BrdU incorporation in neutrophils was analyzed by flow cytometry.

### Single-cell energy metabolism measurements

Single-cell energy metabolism was determined by quantifying protein synthesis in freshly isolated cells subjected to various short metabolic interventions using the flow cytometry-based SCENITH method (Argüello et al., 2020). Briefly, single-cell suspensions were prepared without enzymatic digestion by trituration with a pipette from kidney and liver, and by mincing with scissors or crushing with a pestle and mortar from embryonic and adult bone marrow, respectively. 4×10^5^ cells suspended in RPMI-1640 supplemented with 10% (v/v) FCS and 2 mM L-glutamine (25030-024, Gibco) were seeded into a 96-well plate and treated sequentially with 90 µl of 100 mM 2-deoxy-D-glucose (D6134, Sigma-Aldrich) for 10 min to inhibit glycolysis and/or with 10 µl of 20 µM oligomycin A (75351, Sigma-Aldrich) for 15 min to suppress oxidative phosphorylation. Thereafter, 10 µl of puromycin (0.2 mg/ml, P7255-25MG, Sigma), which incorporates into proteins and terminates protein synthesis, was added to the samples for 30 min at 37 C/5% CO2. Finally, the samples were placed on ice to stop metabolic processes, washed with ice-cold PBS, and stained for flow cytometry. After staining of surface antigens, the cells were permeabilized and stained with a mouse anti-puromycin antibody (clone 12D10, MABE343, Sigma-Aldrich) followed by AlexaFluor Plus 647-conjugated anti-mouse IgG secondary antibody (A32728, Thermo Fisher).

### NETosis assay

Single-cell suspensions of adult bone marrow and E17.5 livers were prepared as described above. To yield a leukocyte-enriched cell suspension, cells were purified through a 33% (v/v) Percoll gradient (25 min, 800 *g*, min. acceleration/deceleration), collected from the bottom of the tube, and washed in cold RPMI-1640 medium. The cell suspension was then plated on 12-well diagnostic slides, cleaned with acetone (1.00014.2500, Merck), isopropanol, and distilled H2O, and rested for 20 min at 37 °C. Next, cells were treated with either 25 nM PMA (P1585, Sigma-Aldrich) or an equal amount of DMSO (D4540, Sigma-Aldrich) and 0.5 µM SYTOX Green (S34860, Thermo Fisher) in medium and incubated for 15 min at 37 °C. Samples were fixed with 4% (v/v) paraformaldehyde (PFA, SC-281692, Santa Cruz Biotechnology) for 5 min at RT, washed with PBS, and mounted with ProLong Gold Antifade Mountant (P36930, Thermo Fisher) mounting medium and 1.5H cover glass. Imaging was performed with a spinning disk confocal microscope (described above) using a Zeiss Plan-Apochromat 63x/1.4 Oil DIC objective and Immersol 518 F (444960-0000-000, Zeiss) immersion oil.

### Phagocytosis assay

Adult bones (femora) were crushed using a pestle and mortar, embryonic bones (femora and tibiae), liver, and kidney were cut into pieces with scissors. Tissues were homogenized into RPMI-1640 supplemented with 10% (v/v) FCS and 2 mM L-glutamine by trituration with a pipette. 5% of recovered cells from an embryonic liver, all cells from two embryonic kidneys, all cells from four embryonic bone marrows (two femora and two tibiae), and 5% of cells from adult bone marrow (one femur) were seeded into a 96-well plate. Cells were plated without leukocyte separation or cell counting to minimize cell losses from minute embryonic tissues. After a 30 min recovery period, 0.5-µm FluoSpheres Carboxylate-Modified Microspheres (1:1000, F8813, Thermo Fisher) were added to the wells, and the samples were incubated at 37°C or on ice (negative controls) for one hour. Thereafter, the cells were collected, washed with ice-cold PBS, and stained for flow cytometry.

### ROS production assay

Single-cell suspensions were prepared by trituration with a pipette in RPMI-1640 supplemented with 10% (v/v) FCS and 2 mM L-glutamine. Thereafter, 3×10^5^ (embryonic liver and adult bone marrow) or 5×10^5^ (embryonic kidney) cells were plated on a 96-well plate. Cells were allowed to rest for one hour at 37 °C or on ice (negative control). CellROX Green reagent (500 nM, C10492, Thermo Fisher) was then added to the samples with or without 50 nM PMA for 20 min at 37 °C or on ice (control). The cells were then collected using a 3 min centrifugation at 500 *g* at 4 °C, washed twice with ice-cold PBS, and stained for flow cytometry.

### Dexamethasone treatment and LPS exposure

For embryonic dexamethasone treatment or LPS exposure experiments, pregnant dams at E17.5 were injected i.p. with either 5 mg/kg of dexamethasone (D2915, Sigma-Aldrich) in PBS or 100 µg/kg LPS (from *E. coli* O55:B5; L4524, Sigma-Aldrich) in 0.9% (w/v) NaCl solution or the equivalent volume of the respective solvent. Animals were euthanized 2 or 4 hours after dexamethasone or LPS treatment, respectively, as described above.

### Antibody treatment experiments

For depletion of embryonic neutrophils, pregnant dams at E12.5, E14.5, and E16.5 were injected i.p. with either 1 mg of anti-Ly-6G (BP0075-1, Bio X Cell) or IgG2a control antibody (BE0089, Bio X Cell). Animals were euthanized at E17.5, and livers were collected and processed for FACS analysis as described before. A secondary anti-rat IgG-AF488 antibody (A11006, Thermo Fisher) staining against the injected anti-Ly-6G antibody was performed before staining with conjugated antibodies for flow cytometry.

For the isotype-switch experiment, pregnant dams were injected i.p. at E12.5 with either 250 µg of anti-Ly-6G antibody or IgG2a control antibody, after which they were injected 100 µg of anti-Ly-6G or control antibody daily until E16.5. Alongside the daily injections, the dams were administered 100 µg of anti-rat kappa IgG light chain isotype (BE0122, Bio X Cell) i.p. every other day at E12.5, E14.5, and E16.5. At E17.5, the animals were euthanized, and the livers were processed for FACS analysis.

### Gfi1 bulk RNA-sequencing

Kidneys (both) and testes (both) from male WT and *Gfi1^mut/mut^* E17.5 embryos (N=3 from pooled samples n=1-2) were collected in RNAlater RNA Stabilization Reagent and stored frozen until processing. RNA was isolated with either the NucleoSpin RNA, Mini kit for RNA purification (kidneys) or RNeasy Plus Micro Kit (testes, 74034, Qiagen) according to the manufacturer’s protocol. Sample quality was assessed using an Agilent Advanced Analytical Fragment Analyzer and Stranded mRNA Library Preparation (1000000124518, Illumina), followed by next-generation sequencing performed by the Finnish Functional Genome Centre (Turku Bioscience Centre, Finland) using an Illumina NovaSeq 6000 SP v1.5 (650–800 M reads/run, 2 lanes).

Fastq files, after automatic adapter trimming, were processed in line with recommendations outlined by HiSAT2 and Stringtie (Pertea et al., 2015; Kim et al.). In brief, quality of the reads was assessed using FASTQC (ver. 0.12.1), reads were trimmed using BBMap (ver. 39.06) and aligned to the GRCm38.84 genome using HiSAT2 (ver. 2.2.1). Reads were then sorted by coordinate and filtered by alignment score using a cutoff of greater than 30 using samtools (ver. 1.19.2) (Danecek et al., 2021). Transcript assembly was done using stringtie (ver. 2.2.1) and the read count matrix was prepared using the prepDE.py3 script provided with HiSAT2. To perform the differential expression analysis, the read count matrix was processed using pyDESeq2 (ver. 0.5.0) with the recommended parameters (Muzellec et al., 2023), and logarithmic fold change shrinkage (Zhu et al., 2019) was applied. Genes with an adjusted p value of less than 0.05 and showing an absolute fold change of over 1.5 were considered significant.

For analysis of transposable elements, low-quality nucleotides and reads were removed from raw data by Trimmomatic (ver. 0.39) (Bolger et al., 2014) and the clean data was mapped to the mouse reference genome (GRCm38.101) using STAR (ver. 2.7) (Dobin et al., 2013). The transposable elements were analyzed by TEtranscripts (ver. 2.2.3) (Jin et al., 2015).

### Immunofluorescence and Conventional Histology

For *Gfi1^mut/mut^* testis section histology, tissues from E17.5 and 7-week-old animals were fixed with 10% formalin (HT501128-4L, Sigma-Aldrich) or 4% PFA at 4 °C overnight. After the fixation, the tissues were rinsed with PBS, dehydrated in an ascending ethanol series, embedded in paraffin, and 5-µm sections were cut.

For immunofluorescence staining, slides were deparaffinized, antigens were retrieved with 0.1 M citrate buffer, pH 6 at 95 °C for 2.5 hours, and nonspecific binding was blocked with 2% (v/v) normal donkey serum (017-000-121, Jackson ImmunoResearch) and 1% (w/v) BSA in PBS (blocking buffer).

Next, samples were stained with a primary antibody against LINE-1 ORF1p (Data S5) overnight at 4 °C, followed by a secondary antibody diluted in blocking buffer for 1 hour at RT, and counterstained with 1 µg/mL DAPI (D1306, Thermo Fisher) in PBS for 10 min at RT. Finally, slides were mounted with ProLong Gold Antifade Mountant mounting medium and 1.5H cover glass. Images were acquired using a Leica THUNDER Imager Live Cell equipped with a Leica LED8 light source, DFT51010 filter cube, 20x/0.70 IMM UV objective, and K8 camera operated using LAS X.

For hematoxylin-eosin staining, slides were deparaffinized and stained with Mayer’s hematoxylin (MHS16-500ML, Sigma-Aldrich) and eosin (180072, Reagena). The sections were imaged using a 3DHISTECH Pannoramic 1000 digital scanner with a 40x/0.95 objective and images were rendered with 3DHISTECH CaseViewer (ver. 2.3).

### Statistics

Statistical analysis was performed using GraphPad Prism (ver. 10.0+). All data are presented as mean or geometric mean values ± SD, geometric SD, or SEM as indicated in the figure legend. Statistical significances between groups were determined using the Mann-Whitney test, One-way or Two-way ANOVA with Tukey’s, Dunnett’s, or Šídák’s multiple comparisons test, or Mixed-effects analysis with Tukey’s multiple comparisons test, depending on the data, and are indicated in the figure legend. P-values < 0.05 are considered to be statistically significant.

## Supplemental material

Fig. S1 contains the flow cytometry gating strategy and frequency of neutrophils in newborn tissues, along with neutrophil kinetics and histology in hematopoietic tissues. Fig. S2 shows additional data regarding neutrophil-specific reporter mice and extravascular neutrophils. Fig. S3 contains an expanded analysis of the scRNA-seq dataset. Fig. S4 displays additional data relating to tissue- and age-specific DEGs in neutrophils. Fig.S5 presents analyses of neutrophil responsiveness to immunomodulatory stimuli in embryos. Fig. S6 shows data characterizing neutrophil depletion models.

## Data availability

Raw scRNA-seq data (FASTQ format) are available from the XX repositories of the original depositing authors (XX accessions: xx). Bulk RNA-seq data are available under accession number XX. Source data are available from the corresponding authors upon reasonable request.

## Acknowledgments

We thank Laura Grönfors and Etta-Liisa Virtomaa for their technical assistance. We thank Selina Keppler for providing technical advice on the SCENITH method. We also acknowledge the Cell Imaging and Cytometry Core (CIC), the Single Cell Omics Core, and the Functional Genomics Core at Turku Bioscience (University of Turku and Åbo Akademi University), supported by Biocenter Finland and the Finnish Advanced Microscopy Node of Euro-BioImaging Finland (CIC), for providing services, instrumentation, and expertise.

This study was financially supported by The Research Council of Finland (M. Salmi and P. Rantakari), the Sigrid Juselius Foundation (M. Salmi and P. Rantakari), The Jane and Aatos Erkko Foundation (P. Rantakari), the Lapland Regional Fund of Finnish Cultural Foundation (E. Lokka), and the Turku Doctoral Program of Molecular Medicine (J. Hofmann, L. Lintukorpi, and E. Lokka).

The authors declare no competing financial interests.

## Author contributions

J. Hofmann, L. Lintukorpi, E. Lokka, S. Cisneros Montalvo, V. Ojasalo, H. Gerke performed experiments; J. Hofmann, D. Kaukonen, and L. Ma performed transcriptomic analyses; N. Kotaja, M. Salmi, and P. Rantakari designed and supervised experiments; and M. Salmi and P. Rantakari designed and supervised the study. M. Salmi and P. Rantakari wrote the manuscript, which was edited by all authors.

---

**Figure S1.**
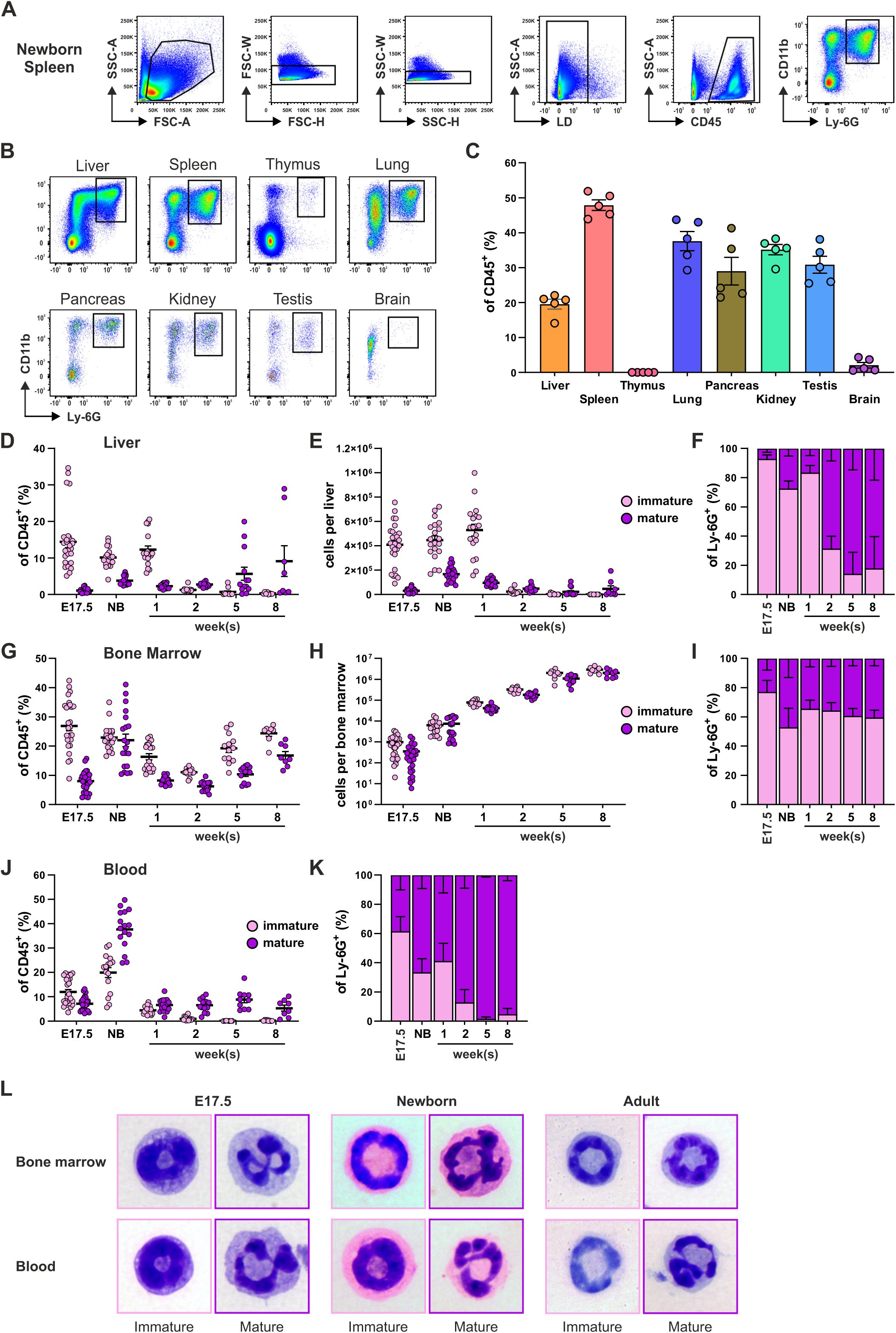
Immature and mature neutrophils are present in steady-state hematopoietic and non-hematopoietic tissues in embryos. **(A)** Flow cytometry gating strategy with representative plots from newborn spleen to identify CD11b^+^Ly-6G^+^ neutrophils. **(B)** Representative plots of Ly-6G^+^ neutrophil population in the indicated newborn tissues. **(C)** Frequency of Ly-6G^+^ neutrophils in the indicated newborn tissues. **(D-K)** Frequency (D, G, J), numbers (E and H), and proportions (F, I, K) of immature CD101*^−^*CXCR2*^−^* and mature CD101^+^CXCR2^+^ neutrophils in the liver (D-F), bone marrow (G-I), and blood (J and K) at the indicated developmental time-points. **(L)** Representative cytological images of sorted Wright-Giemsa-stained immature and mature neutrophils in bone marrow and blood. In C-E, each dot represents one mouse: (C) n = 5 mice (both kidneys and testes were pooled from animal), (D and E) E17.5 n = 27, newborn; n = 20, 1 week; n = 19, 2 weeks; n = 16, 5 weeks; n = 13 and 8 weeks; n = 8 mice. In G and H, each dot represents both femora and tibiae from one mouse (E17.5 n = 29 mice) or both femora of one mouse (newborn; n = 20, 1 week; n = 19 and 2 weeks; n = 16 mice) or one femur from each mouse (5 weeks; n = 13 and 8 weeks; n = 8 mice). In J, each dot represents one mouse (E17.5 n = 27, newborn; n = 16, 1 week; n = 19, 2 weeks; n = 16, 5 weeks; n = 10 and 8 weeks; n = 8 mice). Data are presented as mean ± SEM (C-E, G, H, and J) or ± SD (F, I, and K). Data are from 1 (A-C) or 3-8 (D-K) independent experiments.

**Figure S2.**
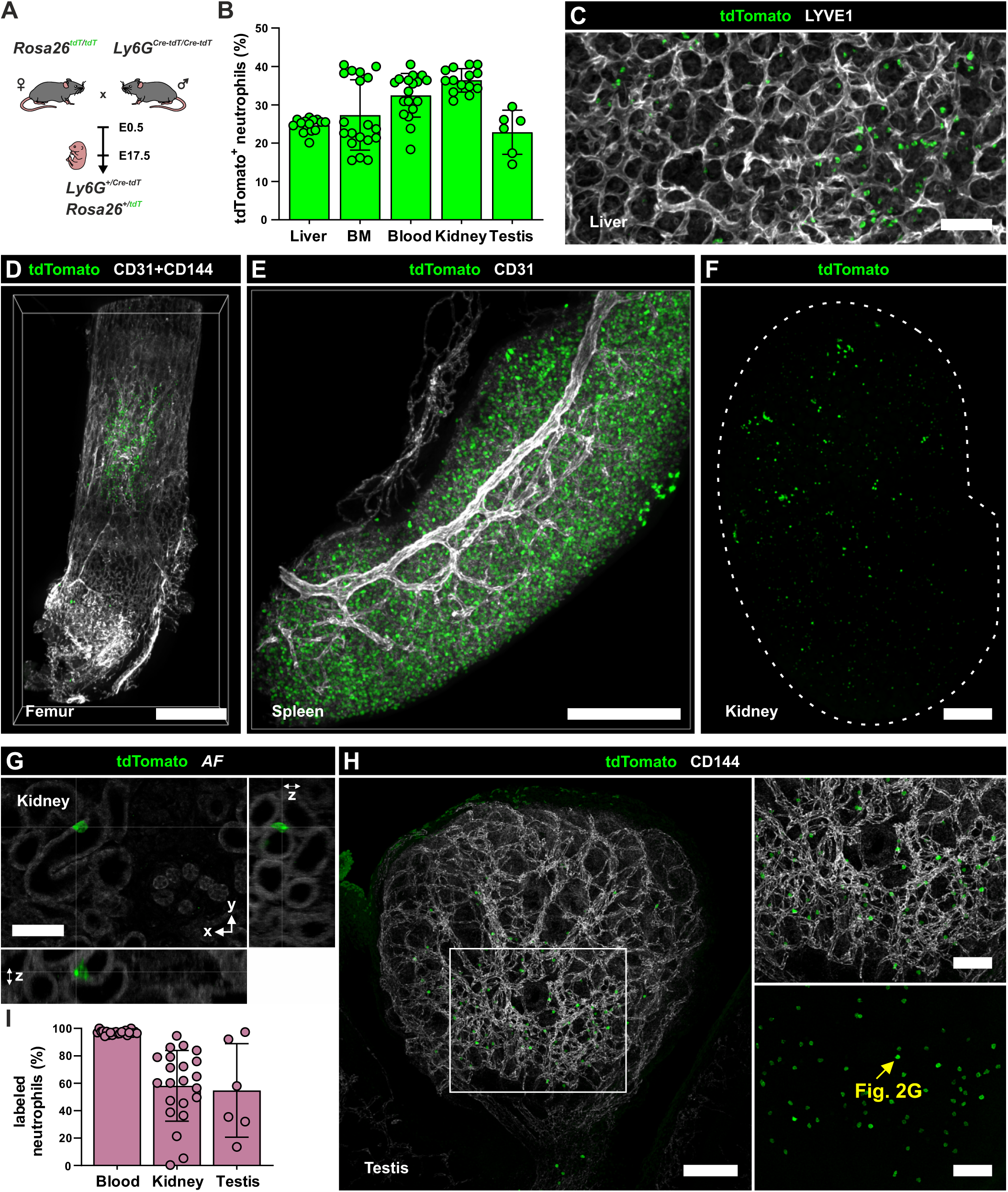
Extravascular neutrophils are abundant in non-hematopoietic and hematopoietic tissues in steady-state embryos. **(A)** Schematic overview of neutrophil reporter crossings used for volumetric imaging of neutrophils in embryonic tissues. **(B)** Frequency of neutrophils labelled by tdTomato expression in indicated tissues of embryonic (E17.5) neutrophil reporter mice. **(C-E)** Representative confocal immunofluorescence microscopy images of optically cleared wholemount liver **(C)**, bone marrow within the femur **(D)**, and spleen **(E)** showing tdTomato-labelled (tdT^+^) neutrophils (green) and vasculature (gray) labelled as indicated from embryonic (E17.5) neutrophil reporter mice. Scale bars 50 µm, 400 µm, and 150 µm, respectively. **(F)** Overview image of the distribution of tdT^+^ neutrophils in the kidney. Scale bar 200 µm. The same image has been used in Fig. 2A. **(G)** Magnified cross-sectional views of tdT^+^ neutrophils localized between the tubule epithelial cells in the kidney. Scale bar 50 µm. **(H)** Volumetric cross-section of testis with connected epididymis and magnified views of the mediastinum and the rete testis area (box) highlighting tdT^+^ cell (arrow) shown in Fig. 2 G. Scale bars 100 µm (full), 50 µm (magnified). Different perspectives of the same images have been used in Fig. 2G. **(I)** Frequency of neutrophils labelled by anti-Ly-6G antibody (clone 1A8) administered intravascularly via umbilical cord in wild-type mice at E17.5. In B and I, each dot represents one mouse, except for testis, where each dot represents a pool of 4 testes from two mice. Data are presented as mean ± SD. In I, data are from 6 independent experiments.

**Figure S3.**
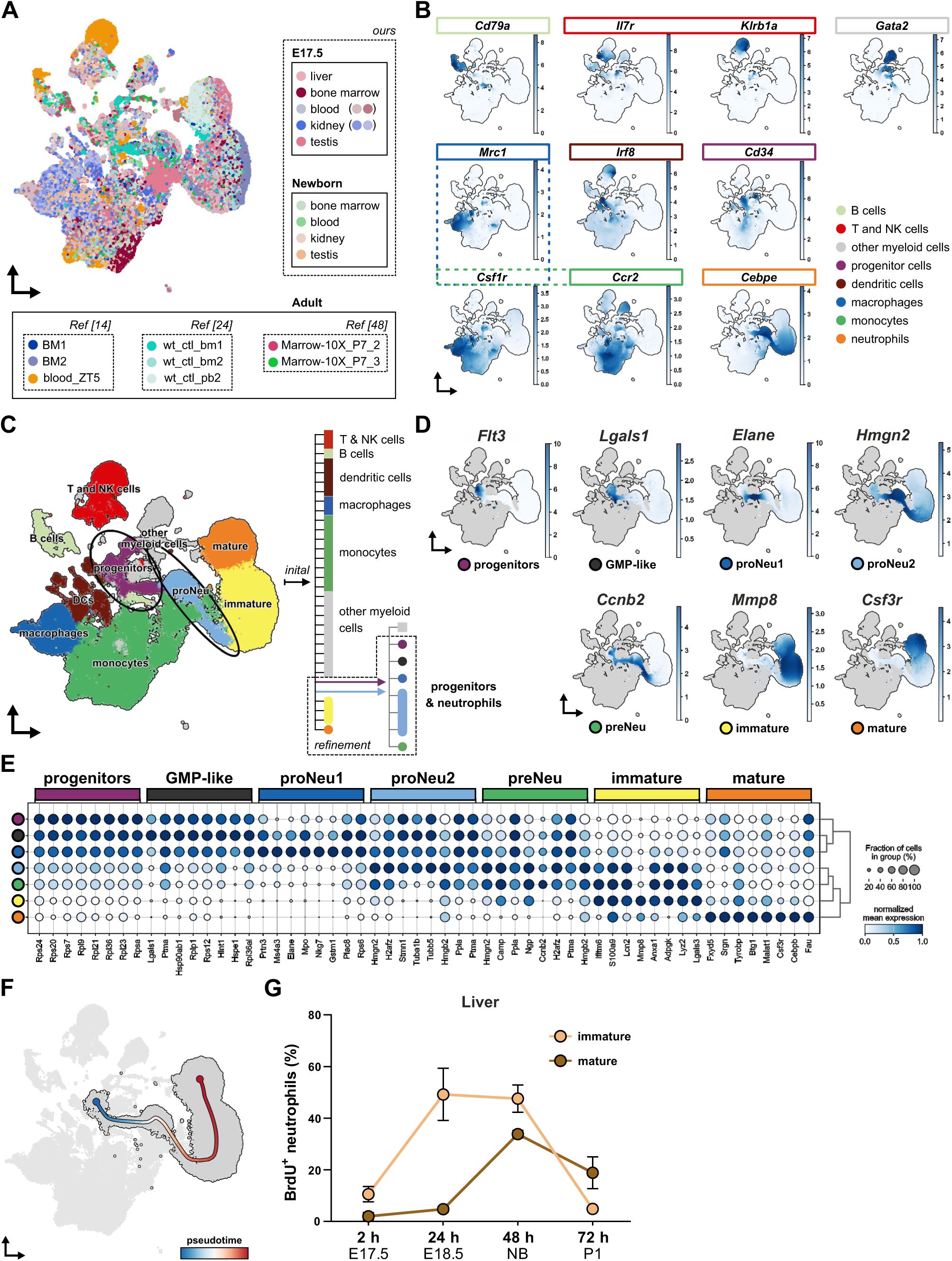
Identification of embryonic neutrophil subpopulations and genes enriched in tissue-resident neutrophils. **(A)** UMAP of our complete integrated scRNA-sequencing dataset highlighting the individual samples from E17.5, newborn, and adult mice **(B)** UMAP-projected marker gene expression for major leukocyte populations using Markov affinity-based graph imputation of cells. **(C)** UMAP and flow graph showing initial cluster annotation (right), annotation refinement strategy (left) for highlighted (black) under-segmented neutrophil progenitor clusters, and final neutrophil cluster annotation shown in Fig. 3B. **(D)** UMAP-projected neutrophil subpopulation marker gene expression using Markov affinity-based graph imputation of cells. **(E)** Dotplot of normalized gene expression for the top 8 ranked (Wilcoxon rank-sum with Bonferroni correction) marker genes across neutrophil subpopulations. **(F)** Neutrophil maturation pseudotime trajectory. **(G)** Kinetics of BrdU incorporation in immature (light) and mature (dark) S-phase neutrophils isolated from embryonic liver at E17.5 after 2, 24, 48, or 72 h BrdU administration to the dam. In G, data are shown as mean with SD, and are from 1-3 experiments per time point.

**Figure S4.**
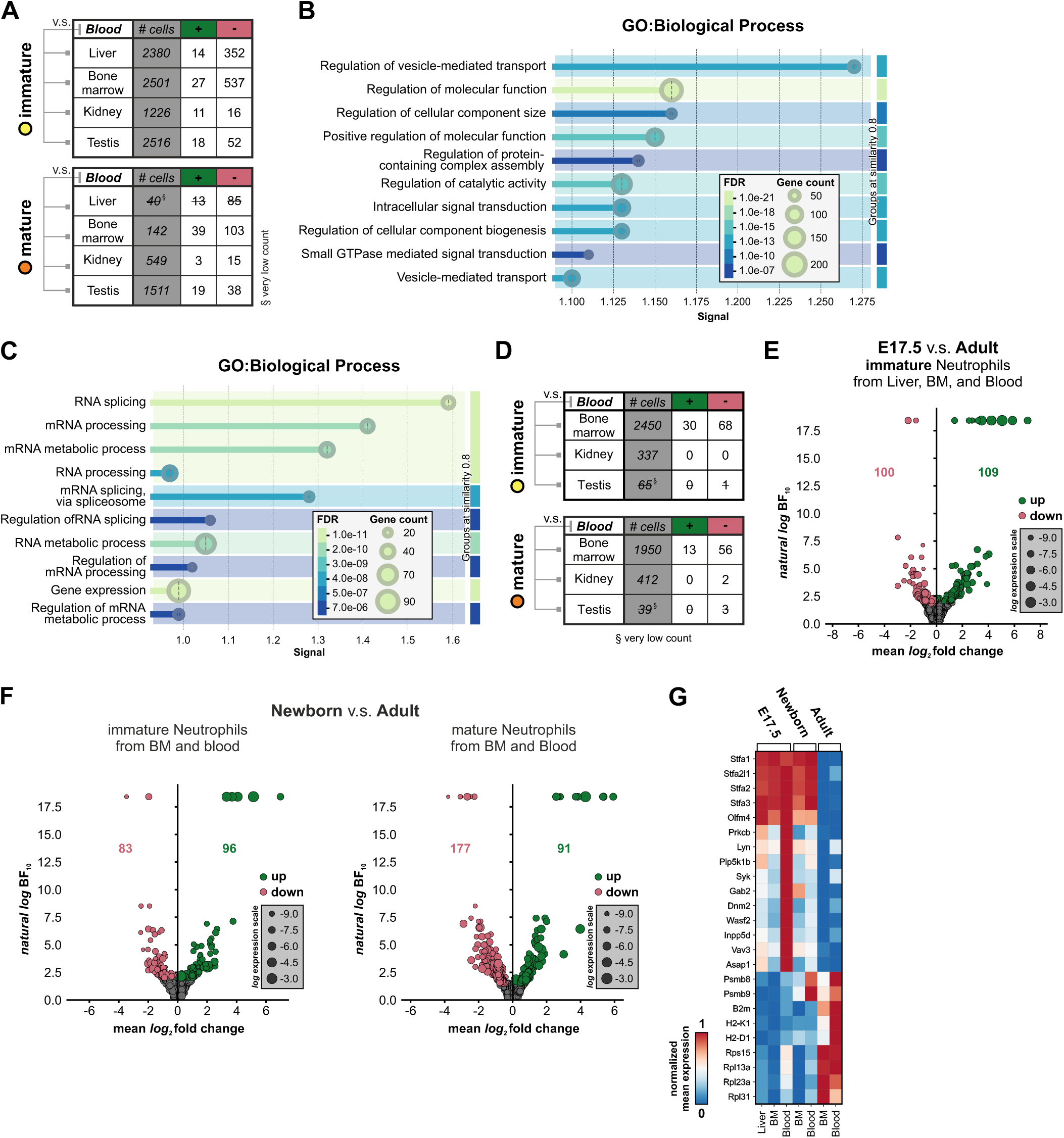
Tissue origin and developmental age distinctly shape neutrophil transcriptomes. **(A)** Number of genes differentially regulated (up, green +; down, red -) in immature (top) or mature (bottom) neutrophils from kidney, testis, liver, and bone marrow compared to cells isolated from blood at E17.5. **(B and C)** STRING functional enrichment visualization of the top 10 grouped gene ontology (GO) terms related to biological processes enriched in genes significantly downregulated in immature neutrophils of bone marrow (B) and liver (C) compared to blood at E17.5. **(D)** Number of genes differentially regulated (up, green +; down, red -) in immature (top) or mature (bottom) neutrophils from kidney, testis, liver, and bone marrow compared to cells isolated from blood in newborns. **(E)** Volcano plot assessing differential gene expression between immature neutrophils isolated from embryonic liver, bone marrow, and blood compared to those in adult bone marrow and blood. **(F)** Volcano plots assessing differential gene expression between immature (left) and mature (right) neutrophils from bone marrow and blood between newborn and adult **(G)** Matrixplot of normalized expression for selected genes differentially expressed in immature neutrophils across ages and tissues. FDR, false discover rate corrected p-values; signal, weighted harmonic mean between the observed/expected ratio and -log(FDR).

**Figure S5.**
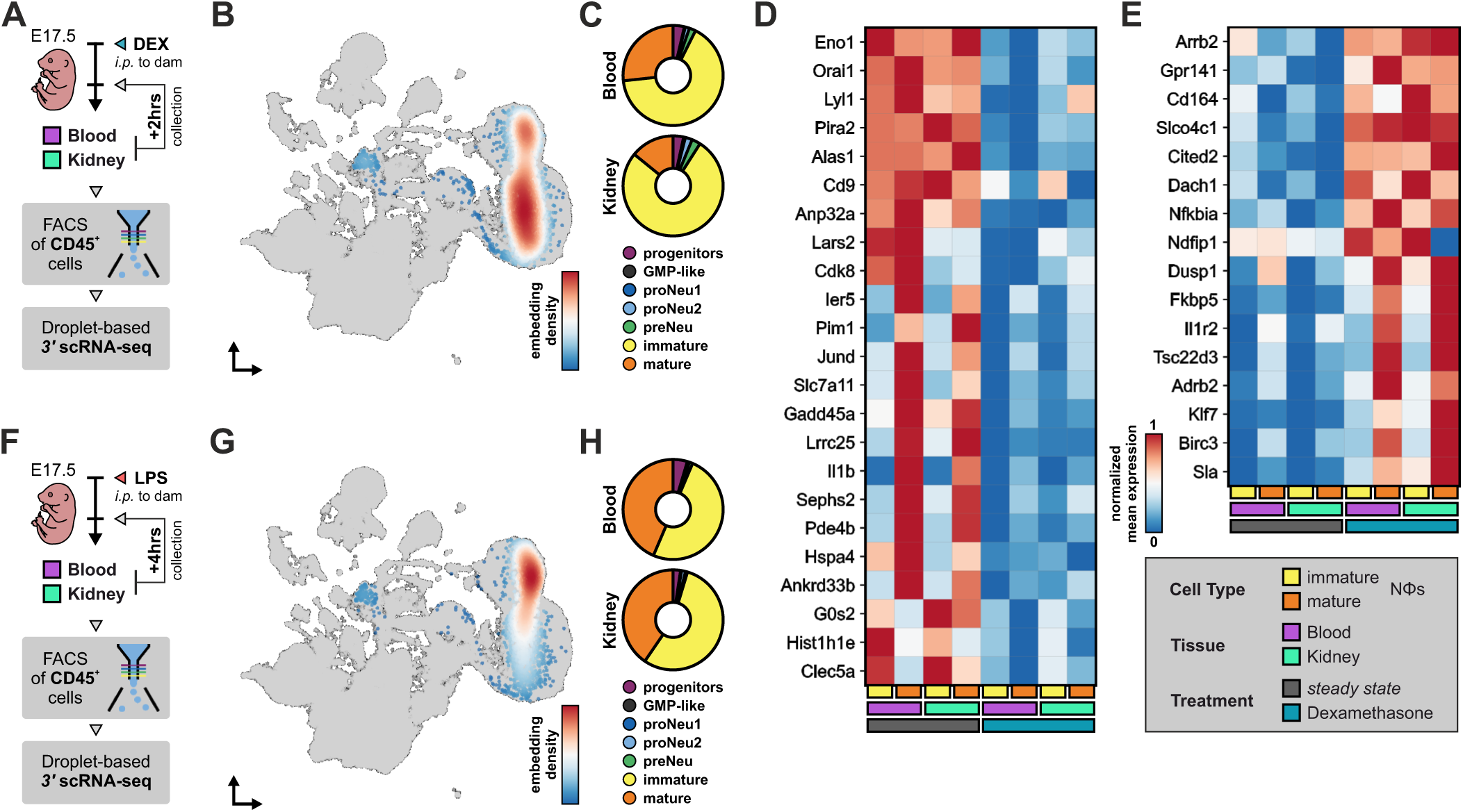
Analyses of neutrophil responsiveness to immunomodulatory stimuli in embryos. **(A)** Schematic outline of dexamethasone treatment and sample collection for scRNA-seq. **(B)** UMAP embedding density (cell distribution heatmap) of neutrophil ontogeny from blood and kidney of E17.5 embryos exposed to dexamethasone treatment. **(C)** Frequency plots of neutrophil subpopulations in blood and kidney of E17.5 embryos exposed to dexamethasone treatment. **(D and E)** Matrixplots of normalized expression for selected genes differentially expressed by consensus (detected in at least 2 of 4 comparisons) in mature and/or immature neutrophils from blood and/or kidney. Genes expression is shown by neutrophil subtype, tissue, and treatment status for genes down- (D) and upregulated (E) in dexamethasone exposed cells. **(F)** Schematic outlining maternal LPS exposure and sample collection for scRNA-seq. **(G)** UMAP embedding density (cell distribution heatmap) of neutrophil ontogeny from blood and kidney of E17.5 embryos after maternal LPS exposure. **(H)** Frequency plots of neutrophil subpopulations in blood and kidney of E17.5 embryos after maternal LPS exposure. DEX, dexamethasone; i.p., intraperitoneal; FACS, fluorescence-activated cell sorting; NΦs, neutrophils; LPS, lipopolysaccharide.

**Figure S6.**
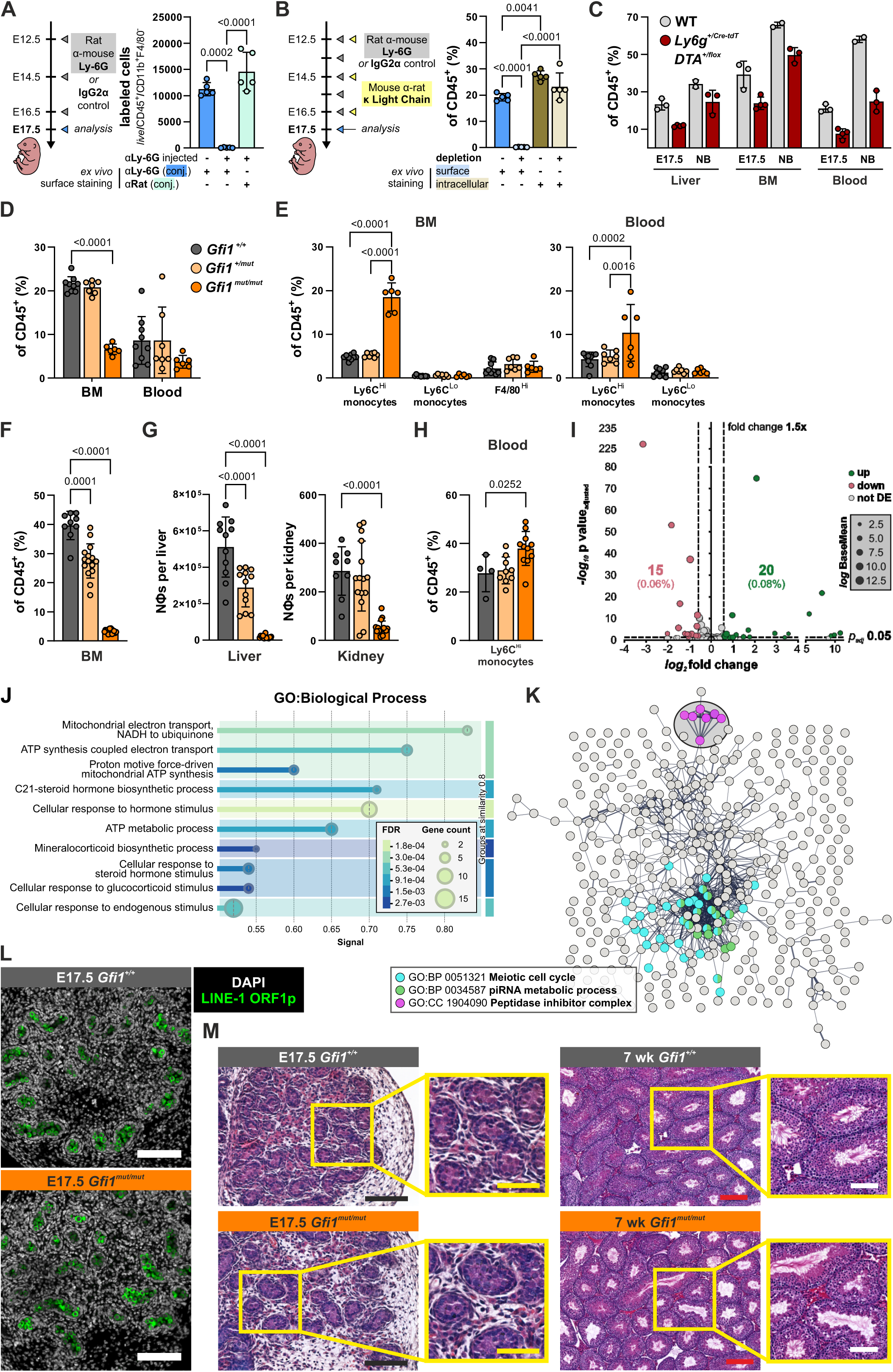
Genetic neutrophil depletion models and their effects on testes. (A and. **B)** Schematic overview and flow cytometric analysis of (A) α-Ly-6G antibody-mediated and (B) isotype-switch neutrophil depletion in embryonic liver. **(C)** Frequency of Ly-6G^+^ neutrophils in indicated tissues isolated from E17.5 and newborn *Ly6g^+/Cre-tdT^* x DTA^+/flox^ mice. **(D and E)** Frequency of Ly-6G^+^ neutrophils (D) and Ly-6C^Hi^ and Ly-6C^Lo^ monocytes and F4/80^Hi^ macrophages (E) in bone marrow and blood of 4-wk-old *Gfi1^+/+^* (gray), *Gfi1^+/mut^* (light orange), and *Gfi1^mut/mut^* (orange) mice. **(F)** Frequency of Ly-6G^+^ neutrophils in bone marrow of E17.5 *Gfi1^+/+^*, *Gfi1^+/mut^*, and *Gfi1^mut/mut^* mice. **(G)** Number of Ly-6G^+^ neutrophils in E17.5 liver (left) and kidney (right) of E17.5 *Gfi1^+/+^*, *Gfi1^+/mut^*, and *Gfi1^mut/mut^* mice. **(H)** Frequency of Ly-6C^Hi^ monocytes in the blood of E17.5 *Gfi1^+/+^, Gfi1^+/mut^, and Gfi1^mut/mut^ mice*. **(I)** Volcano plot of bulk RNA-sequencing assessing differential gene expression in kidneys between *Gfi1^mut/mut^* and WT embryos at E17.5. **(J)** STRING functional enrichment visualization of the top 10 grouped gene ontology terms related to biological processes enriched in genes significantly upregulated in the testes of *Gfi1^mut/mut^* compared to WT embryos at E17.5. **(K)** STRING network of genes (nodes) significantly downregulated in testes of *Gfi1^mut/mut^* compared to WT embryos at E17.5, highlighting genes associated with selected gene ontology terms. Line thickness indicates the degree of confidence of the predicted interaction. **(L)** Representative widefield immunofluorescence microscopy images of testis sections from *Gfi1^+/+^*(left) and *Gfi1^mut/mut^* (right) E17.5 embryos stained for LINE-1 open reading frame 1 protein (ORF1p; green) and DNA (gray). Scale bars 100 µm. **(M)** Representative HE-staining of testis morphology in E17.5 (left) and 7-week (right) *Gfi1^+/+^* (top) and *Gfi1^mut/mut^* (bottom) embryos/mice. Scale bars 100 µm (black and white), 200 µm (red), and 50 µm (yellow). In A-H, mean with SD are shown. (A, B, and E) Two-way ANOVA/Tukey’s multiple comparisons test, (D and F) Two-way ANOVA/Dunnett’s multiple comparisons test, (G and H) One-way ANOVA/Dunnett’s multiple comparisons test. In A-C, data are from one experiment, in D-G, 4-5 independent experiments, and in H, from 3 independent experiments. FDR, false discover rate corrected p-values; signal, weighted harmonic mean between the observed/expected ratio and - log(FDR).

